# Zebrafish Neuromesodermal Progenitors Undergo a Critical State Transition *in vivo*

**DOI:** 10.1101/2022.02.25.481986

**Authors:** Kane Toh, Dillan Saunders, Berta Verd, Benjamin Steventon

## Abstract

The transition state model of cell differentiation proposes that a transient window of gene expression stochasticity precedes entry into a differentiated state. As this has been assessed primarily *in vitro*, we sought to explore whether it can also be observed *in vivo*. Zebrafish neuromesodermal progenitors (NMps) differentiate into spinal cord and paraxial mesoderm at the late somitogenesis stages. We observed an increase in gene expression variability at the 24 somite stage (24ss) prior to their differentiation. From our analysis of a published 18ss scRNA-seq dataset, we showed that the NMp population possesses a signature consistent with a population undergoing a critical transition. By building *in silico* composite gene expression maps from our image data, we were able to assign an ‘NM index’ to each *in silico* NMp based on the cumulative expression of its neural and mesodermal markers. With the NM index distributions, we demonstrated that cell population heterogeneity of the NMps peaked at 24ss. We then incorporated stochasticity and non-autonomy into a genetic toggle switch model and uncovered the existence of rebellious cells, which we then confirmed by reexamining the composite maps. Taken together, our work supports the transition state model within an endogenous cell fate decision making event.

## Introduction

Neuromesodermal progenitors (NMps) are axial progenitors that co-express the lineage-specific transcription factors *Brachyury/T/Tbxta* and *Sox2* and are competent to generate both neural (e.g. spinal cord) and mesodermal (e.g. somite) fates at the single-cell level (Henrique et al., 2015; Wymeersch et al., 2021). Bipotent NM cells have been identified in amniotes such as mouse (Cambray & Wilson, 2007; Cambray & Wilson, 2002) and chick (Brown & Storey, 2000; Guillot et al., 2021; Wood et al., 2019) as well as anamniotes such as *Xenopus* (Davis & Kirschner, 2000; Gont et al., 1993), axolotl (Taniguchi et al., 2017) and zebrafish (Martin & Kimelman, 2012). Therefore, they are an evolutionary conserved cell population whose decision to generate spinal cord and paraxial mesoderm provides an ideal system to explore the mechanisms of cell fate decision making *in vivo*.

The degree to which NMps divide to produce daughter cells of both neural and mesodermal fates depends on species-specific growth dynamics (Steventon & Martinez-Arias, 2017). In the zebrafish embryo, there is little volumetric growth associated with posterior body elongation (Steventon et al., 2016) and proliferation stops abruptly within the embryo around the 10 somite stage (ss) (Bouldin et al., 2014; Zhang et al., 2008). Correspondingly, zebrafish tailbud NMps are a largely quiescent pool of monofated progenitors that give rise to a limited portion of the posterior body axis (Attardi et al., 2018; Bouldin et al., 2014). In the mouse embryo, using retrospective clonal analysis, long clones originating from a single cell have been observed in both neural and mesodermal tissues (Tzouanacou et al., 2009), which is consistent with a proliferative phase in the mouse NMps at around E9.5 (Wymeersch et al., 2016). Despite this difference in developmental dynamics, two independent lines of evidence support the notion that zebrafish NMps are, like all other vertebrate NMps, competent towards both neural and mesodermal fates. Firstly, single cell transplantation experiments demonstrate that zebrafish NMps can be steered towards either neural or mesodermal fates upon manipulation of the canonical Wnt pathway (Martin & Kimelman, 2012). Secondly, a single cell transcriptomic signature that contains conserved markers of both spinal cord and paraxial mesoderm states have been discovered for the zebrafish NMps at late gastrulation/early tailbud stages of development (Lukoseviciute et al., 2021). Thus, a conceptual clarification between NM competent cells and NM progenitors (NMps) has been proposed, of which a differing proportion of NM competent cells act as NMps in a stage and specific-specific manner dependent on the rate of proliferation (Binagui-Casas et al., 2021; Sambasivan & Steventon, 2021). In this paper, we refer to these cells as zebrafish tailbud ‘NMps’ to remain consistent with previous literature, although they are better understood as NM competent cells at post 10ss of development.

How do these zebrafish tailbud NMps differentiate into their NM derivatives? Differentiation has been widely characterized as an ordered and largely deterministic succession of cellular states, specifically transcriptomic states, that emerge from the activation of a set of master transcription factors in a gene regulatory network (Davis et al., 1987; Whyte et al., 2013). If transcriptomic states strongly correlate to developmental lineage, then we can sort single cells along a pseudotemporal axis of developmental progression using their transcriptomic states as the similarity measure and infer the gene expression trajectories within these differentiating cells. Elucidating the pseudotemporal axis has uncovered numerous insights into development (Wagner et al., 2018; Wolf et al., 2019) and disease (Mukherjee et al., 2020; Petti et al., 2022). Despite their utility, pseudotemporal ordering algorithms make a critical simplifying assumption: cells with similar transcriptomic profiles are assigned to be closer together in their developmental maturity along a lineage (Schier, 2020; Tritschler et al., 2019). This biological assumption has been challenged by several observations. First, *in vitro* studies revealed the prevalence of non-genetic heterogeneities within clonal stem cell populations, where cells stochastically transition between distinct metastable states despite being functionally homogeneous (Canham et al., 2010; Hayashi et al., 2008; Trott et al., 2012). In addition, global transcriptomic trajectories may be driven by complex dynamics such as slow fluctuations that persist across cell division cycles (Chang et al., 2008) and oscillatory dynamics in key regulators (Verd et al., 2018). Furthermore, distinct trajectories may converge to the same terminal fate (Packer et al., 2019). These observations suggest that the relationship between cell fate and transcriptomic state can be complex (Casey et al., 2020) and additional information is required before constraining the possible dynamics that arise from snapshot data with the maximum parsimony assumption (Tanay & Regev, 2018; Weinreb et al., 2018).

An alternative class of models proposes that differentiation is a two-stage process: an initial regime of increased heterogeneity within the differentiating population due to elevated stochasticity in gene expression, followed by the stabilisation of the gene expression pattern during cell-fate determination. During the initial stochastic phase, transcriptomic states and cell fates are less correlated as gene expression heterogeneity increases. Models that belong in this class include the Darwinian model of cellular differentiation (Kupiec, 1997; Minelli et al., 2014; Paldi, 2020), the ‘exploratory’ model of stem cell decision-making (Halley et al., 2009) and the ‘transition state’ model (Antolović et al., 2019; Arias & Hayward, 2006; Brackston et al., 2018; Moris et al., 2016; Muñoz-Descalzo et al., 2012; Rué & Martinez Arias, 2015). Taking a statistical mechanical perspective, this phenomenon of ‘regulated stochasticity’ is consistent with cellular differentiation being a critical phase transition (Teschendorff & Feinberg, 2021). Experimental observations of a surge in gene expression variability that precedes a commitment phase are found predominantly in *in vitro* models such as hematopoietic stem cell differentiation models (Hu et al., 1997; Mojtahedi et al., 2016; Moussy et al., 2017; Pina et al., 2012; Richard et al., 2016), induced pluripotent stem cells (iPSCs) (Bargaje et al., 2017; Buganim et al., 2012) and mouse embryonic stem cells (mESCs) (Moris et al., 2018; Semrau et al., 2017; Stumpf et al., 2017). In contrast, *in vivo* observations of this phenomenon has been comparatively rare (Antolović et al., 2019; Peláez et al., 2015). *In vivo* evidence are vital to ensure that the preceding *in vitro* observations are not due to artifacts of cell culture conditions (MacArthur & Lemischka, 2013; Smith, 2013) or reporter dynamics (Smith et al., 2017).

In this paper, we assessed the transition state hypothesis *in vivo* during the zebrafish tailbud NMp differentiation event. Our results can be grouped according to two features of the hypothesis:

### 1. Transient increase in transcriptional heterogeneity during NMp differentiation

As photolabels of the NMp region at the 12ss revealed that cells only contribute to somites and spinal cord from the 24 somite level onwards (Attardi et al., 2018), we focused on a time-window between the 18ss and 30ss to capture the commitment event. By quantifying the single-cell levels of nuclear *sox2* and *tbxta* expression in NMps from 18ss to 30ss *in situ*, we demonstrate that the heterogeneity in expression of both genes as well as the variability in NMp number peak at 24ss. In addition, by examining a publicly available 18ss scRNA-seq dataset of the zebrafish embryo (Wagner et al., 2018), we found that NMps have a higher critical index and transcriptional noise relative to their derivatives, supporting the view that the NMp population is undergoing a critical transition. Furthermore, by combining the expression of multiple NMp marker genes across multiple samples with an image alignment pipeline (ZebReg) and computing the ‘NM index’, we found that the Shannon entropy, a measure of the population heterogeneity, also peaks at 24ss.

### 2. Loosening of the relationship between cell state and cell fate: existence of ‘Rebellious’ cells

Given the importance of stochasticity in the initial stage of the transition state model, we modelled the behaviour of a stochastic non-autonomous toggle switch to mimic the gene expression heterogeneity observed during the NMp differentiation event. The incorporation of non-autonomy in the stochastic model reflects the fact that embryonic development *in vivo* is a dynamic process that unfolds over time, where intrinsic changes in a cell’s molecular state exists in a causal feedback loop with extrinsic changes that occur in its surrounding environment (Busby & Steventon, 2021). Interestingly, we identified a subset of trajectories that transiently reside in the competing attractor before eventually switching into the primary attractor at the end of the simulation. Following the work of Mojtahedi and colleagues, we labelled these trajectories as ‘Rebellious’ (Mojtahedi et al., 2016). We exploited the relative biological simplicity of the zebrafish NMp system to relate cellular states to cellular fates by examining the spatial locations of the NMps within our ZebReg composite maps. Reexamining our composite maps, we identified an increase in the number of Rebellious cells at the 24ss within the mesoderm-fated domain.

Taken together, our work supports the existence of a transition state and the presence of ‘rebellious’ cells *in vivo* during zebrafish NMp differentiation.

## Results

### Heterogeneity in *sox2* and *tbxta* expression and variability in the number and locations of NMps peak at 24ss

To assess the number and location of zebrafish tailbud NMps over time, we performed HCR stains for *sox2* and *tbxta* to quantify the mRNA expression of single cells *in situ* within the zebrafish tailbud (Figure S2A-E). First, we compared the expression of nuclear *sox2* and *tbxta* in the NMps against the posterior notochord and posterior neural tube populations (Figure S2F-H’). We find that the posterior notochord population has a tight distribution of nuclear *sox2* with a mean normalised intensity close to 0 and a broader nuclear *tbxta* distribution (Figure 1D-D’). Conversely, the posterior neural tube population has a tight distribution of nuclear *tbxta* with a mean normalised intensity close to 0 and a broader nuclear *sox2* distribution (Figure 1E-E’). On the other hand, the NMps are distinct from both populations as they have broad marginal distributions of both nuclear *sox2* and *tbxta* (Figure 1F-F’’).

**Figure 1.**
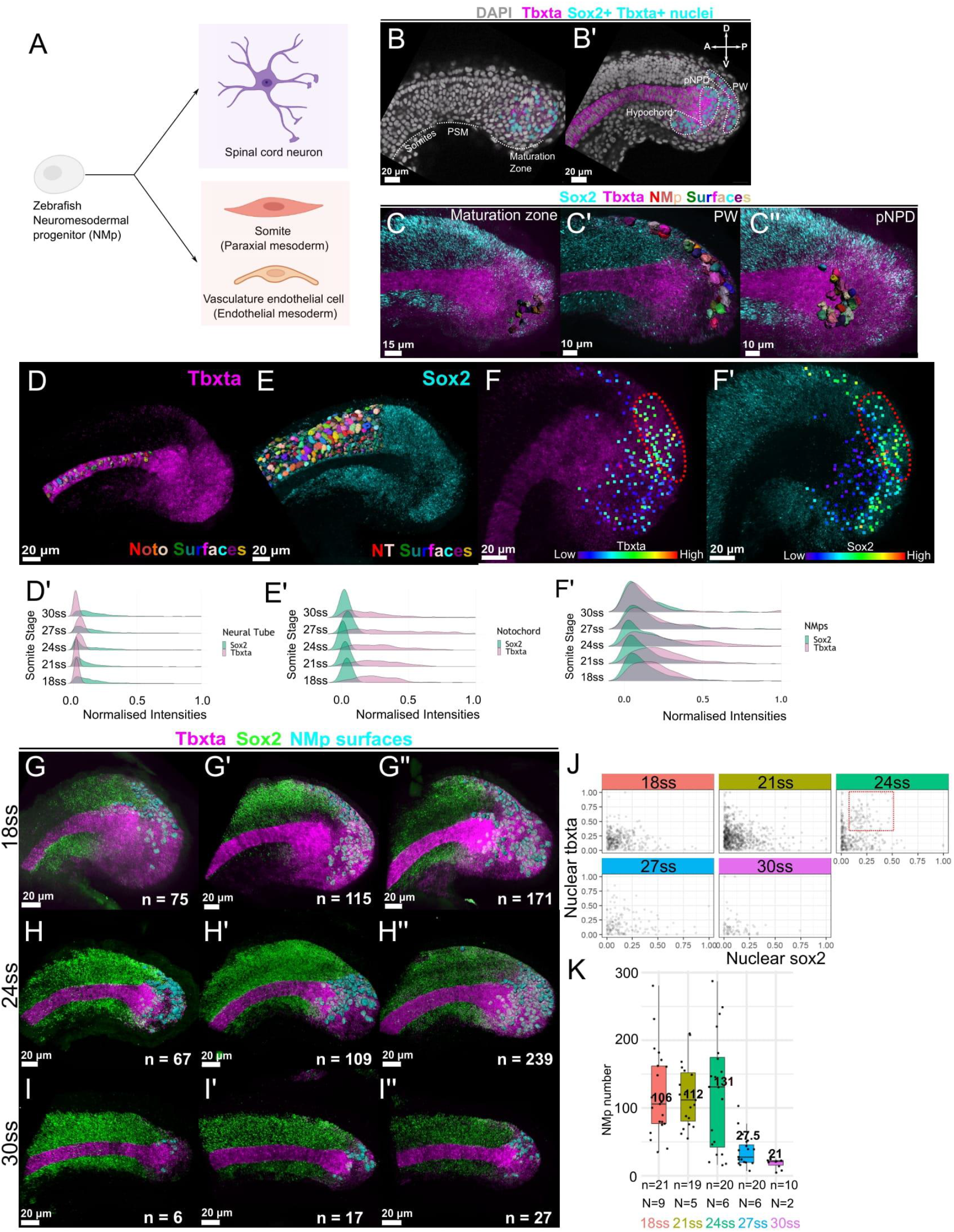
Heterogeneity in *sox2* and *tbxta* expression and variability in the number and locations of NMps peak at 24ss. (A) Zebrafish NMps undertake a binary fate decision to differentiate into the posterior neural and mesodermal fates. (B) 2D lateral slice showing *sox2*+*tbxta*+ nuclei (cyan surfaces) in the maturation zone. (B’) 2D medial slice showing *sox2*+*tbxta*+ nuclei in the hypochord, pNPD and PW. pNPD: posterior notochord progenitor domain; PW: posterior wall. (C-C’’) Segmented NMp surfaces located in the (C) maturation zone (C’) PW (C’’) pNPD. (D, E, F-F’) Maximum intensity projections of *tbxta* and *sox2* shown alongside segmented surfaces of the posterior notochord (D) and posterior neural tube (E). NMps are shown as points colored according to their (F) *tbxta* and (F’) *sox2* expression levels. The red regions highlight the NMps in the posterior wall that co-express intermediate levels of (F’) *tbxta* and (F’’) *sox2*, which is also highlighted with a red region in (J). (D’, E’, F’’) Ridge plots from 18ss to 30ss depicting the expression distributions of nuclear *sox2* and *tbxta* distributions in the (D’) posterior notochord (E’) posterior neural tube (F’) NMp populations. (G-I’’) HCR-stained samples at (G-G’’) 18ss, (H-H’’) 24ss and (I-I’’) 30ss with three representative images per set. n: number of segmented NMps in each sample. (J) Scatterplots of *sox2* and *tbxta* expression of NMps from 18ss to 30ss at three-somite intervals. Each point corresponds to the normalised nuclear *sox2* and *tbxta* intensities of a single NMp. The red region at 24ss highlights the NMps with intermediate levels of both genes. (K) Box and whisker plots of the number of NMps from 18ss to 30ss at three-somite intervals. Each point corresponds to the number of NMps in a single sample. The median NMp number is indicated in bold. n: total number of samples analysed for each stage (biological replicate). N: number of distinct imaging experiments, where different biological samples imaged on the same day are considered a single imaging experiment.

Next, we quantified the nuclear *sox2* and *tbxta* levels within the NMp population across the different somitogenesis stages. In the gene expression scatterplots (Figure 1J), we find that most NMps are *sox2*+_low_*tbxta*+_low_. However, at 24ss, we also find a greater number of *sox2*+_int_*tbxta*+_int_ NMps, reflecting a transient increase in the transcriptional heterogeneity of the NM gene expression states. We then quantified the number and position of NMps at each stage across multiple individual tailbud samples. We found significant variation in the position (Figure 1G-I) and number (Figure 1K) of NMps across samples at all stages under study. Notably, peak variability in NMp number occurred at the 24ss (Figure 1K). Taken together, our analysis demonstrates that a transient phase of increased heterogeneity in *sox2* and *tbxta* expression states occurs around 24ss. This closely matches the developmental stage at which labelled NMps contribute to both spinal cord and paraxial mesoderm (Attardi et al., 2018) and therefore suggests that the increased heterogeneity precedes the commitment to either NM fate.

### Analysis of 18ss scRNA-seq data reveals a peak in the critical index and transcriptional noise index in the NMp population relative to its derivatives

A second prediction of the transition state model is that cells should explore a larger region of gene expression space prior to cell fate commitment as the progenitor basin flattens, resulting in a more dispersed ‘cloud’ of points in state space (Huang, 2009). Consequently, cell population heterogeneity increases, whereas between-gene variation decreases as cells up-regulate groups of either neural or mesodermal genes in coordinated fashion (Mojtahedi et al., 2016). To assess this in the context of zebrafish NMps *in vivo*, we made use of a recently published single-cell RNAseq dataset at 18ss (Wagner et al., 2018).

First, we reanalysed the scRNA-seq data using an independent dimensional reduction and clustering approach to obtain the 8 tailbud subclusters that include the NMps and their derivatives (See STAR Methods). We include the expression of selected differentially expressed genes for the neural, mesodermal and NMp clusters as dot plots (Figure 2C-C’’) and provide information on marker gene expression for all 8 clusters in Table S3. In support of our manual annotation of the NMp cluster, we find that most of the *sox2*+*tbxta*+ co-expressing cells are found within the NMp cluster, and the NMp cluster is sandwiched between two neural clusters and five mesodermal clusters (Figure 2A-B). This is consistent with *sox2* and *tbxta* emerging as differentially expressed genes in this cluster (Table S3). In addition, we validated a subset of the identified NMp marker genes experimentally via HCR and find that they are all expressed within the NMps within the tailbud (Figure S6), supporting the robustness of our *in silico* analysis.

**Figure 2.**
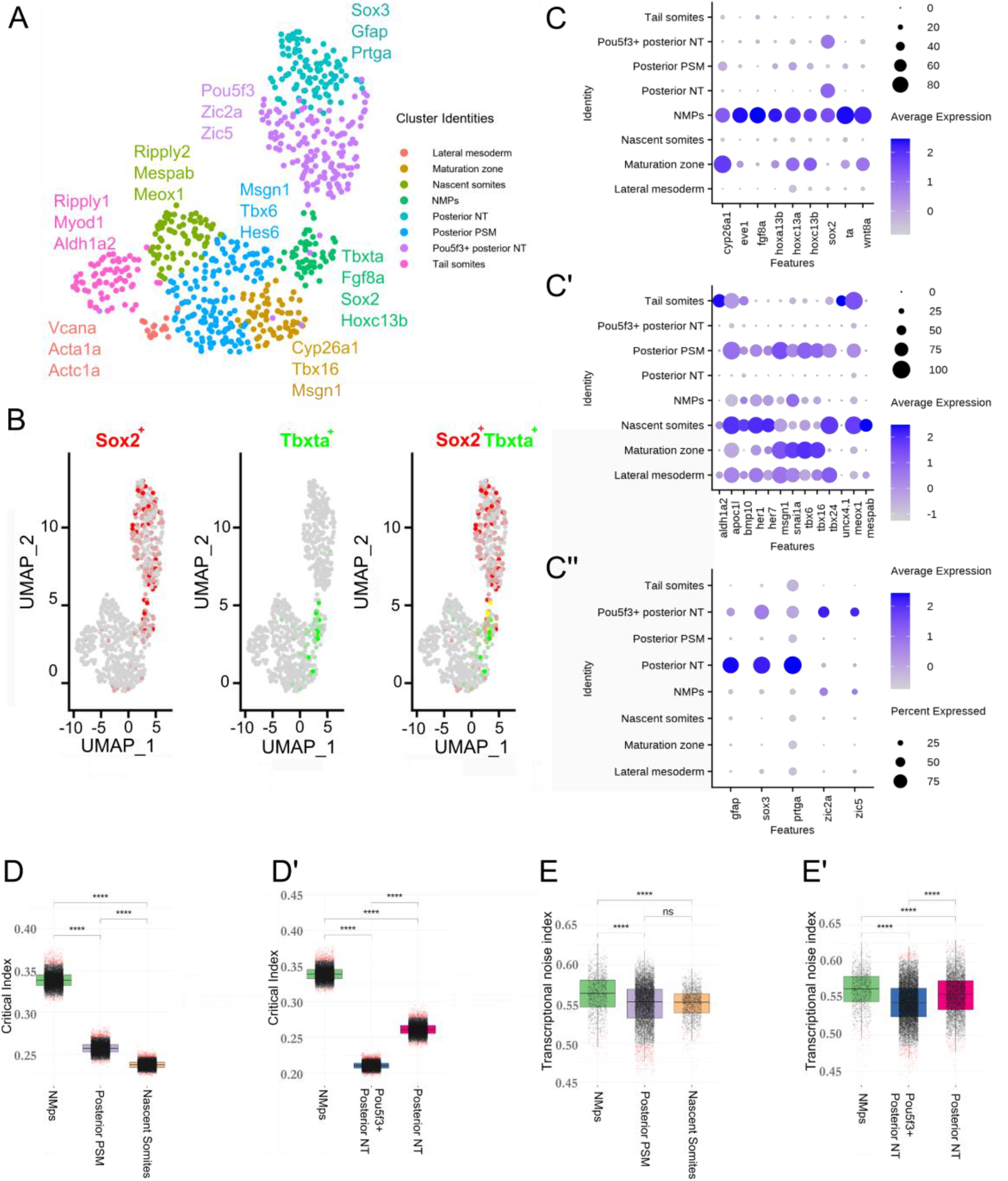
Analysis of 18ss scRNA-seq data reveals a peak in the critical index and transcriptional noise index in the NMp population relative to its derivatives. (A) UMAP embedding showing the 8 tailbud clusters at 18ss alongside the key differentially expressed genes used for manual annotation. UMAP: uniform manifold approximation and projection. (B) UMAP embedding in (A) coloured by *sox2* and *tbxta* expression and *sox2*-*tbxta* together to illustrate co-expression. (C-C’’) Dot plots displaying the expression of differentially expressed genes from the tailbud (C) NMp cluster, (C’) Mesodermal clusters and (C’’) Neural clusters. (D-D’) Distribution of the critical transition index I_c_ calculated using marker genes of each cluster along the mesodermal (D) and neural (D’) branches. A bootstrapping procedure was applied in calculating I_c_ to account for the differences in cell number between cell clusters. (E-E’) Distribution of pairwise cell-to-cell distances / transcriptional noise along the mesodermal (E) and neural (E’) branches.

We find that the NMp cluster is enriched for the posterior Hox genes *hoxc13b, hoxc13a* and *hoxa13b*, with avg_log2FC of 1.02, 0.94 and 0.78 respectively (Table S3). Avg_log2FC measures the log fold-change of the average expression of these genes in the NMp cluster versus the other clusters (Stuart et al., 2019). This is consistent with reports of their roles in mediating the termination of axial elongation (Aires et al., 2019; Young et al., 2009) and directing mesoderm formation in the NMp niche (Ye & Kimelman, 2020). In terms of signalling pathways, *wnt8a* and *fgf8a* appear as the top two genes enriched in the NMp cluster, both of which are actively involved in NMp maintenance and differentiation (Goto et al., 2017; Row et al., 2016). Notably, *fgf8a* is expressed in >80% of cells in the NMp cluster (PCT1 = 0.816) and <8% of cells in all other clusters (PCT2 = 0.075). Finally, our analysis also identified *cyp26a1*, a retinoic-acid degrading enzyme that safeguards the Wnt/*tbxta* positive feedback loop, with an avg_log2FC of 1.02 (Martin & Kimelman, 2010).

Besides identifying the known molecular players in NMp differentiation, we also uncovered numerous other genes involved in a diverse range of processes. Genes with annotations that implicate their roles in signalling pathways feature prominently and include *wls, wnt8-2*, and *depdc7* for Wnt, *angptl2b* and *her12* for Notch, *nog2* and *id3* for BMP and *fgf4* for FGF signalling. Also, three genes were annotated with cytoskeleton-associated processes (*tagln3b* and *enc1* have actin-binding activity and *kif26ab* regulates microtubule motor activity), two possess histone deacetylase binding activity (*znf703* and *kdm6a*) and another two are associated with ubiquitination (*traf4a* and *ubl3a*). Interestingly, *foxd3*, a neural crest marker, emerged as a candidate that is enriched in the NMp cluster, with an adjusted p-value of 1.92×10^−6^. It is expressed in >20% of cells in the NMp cluster and <4% of cells across all the other 7 clusters (Table S3). This observation confirms the results of recent study that revealed a common transcriptomic signature of the neural crest and NM populations (Lukoseviciute et al., 2021).

Cell fate decision making has been proposed to be a critical transition event, with both these features captured in a single ‘Critical Transition Index’ that has previously been shown to predict a cell fate decision making event within blood progenitors as they commit to either myeloid or erythroid lineages (Mojtahedi et al., 2016). In similar vein, we computed the critical indices for the NMp, neural (pou5f3+ posterior NT and posterior NT) and mesodermal (posterior PSM and tail somites) clusters (Figure 2D-D’). Along both the neural and mesodermal differentiation trajectories, the NMp cluster cells have the highest critical index, which is consistent with a cell population undergoing a dynamical bifurcation. Next, we assessed the level of transcriptional noise in the population. We observed that the NMp cluster cells have a higher transcriptional noise relative to the other cell populations (Figure 2E-E’). Therefore, both quantitative indices support the hypothesis that NMps are approaching a critical transition at 18ss.

### Gene expression imputation and the construction of a composite map via ZebReg demonstrates a peak in the NM index entropy at 24ss

Our observation that the number and position of NMps vary extensively between stage-matched embryos, especially at the 24ss, suggests that there is significant variability in *sox2* and *tbxta* expression within the NMps. Consequently, fixed measurements of gene expression from a single sample alone would be inaccurate as it can only give an instantaneous snapshot capturing one out of many different gene expression states that the NMp population can potentially explore. To leverage the gene expression information across multiple tailbud samples, we developed a tool called ZebReg that takes images of stage-matched zebrafish tailbud samples as inputs, converts them into point clouds and registers the point clouds together to construct composite gene expression target maps (Figure 3A).

**Figure 3.**
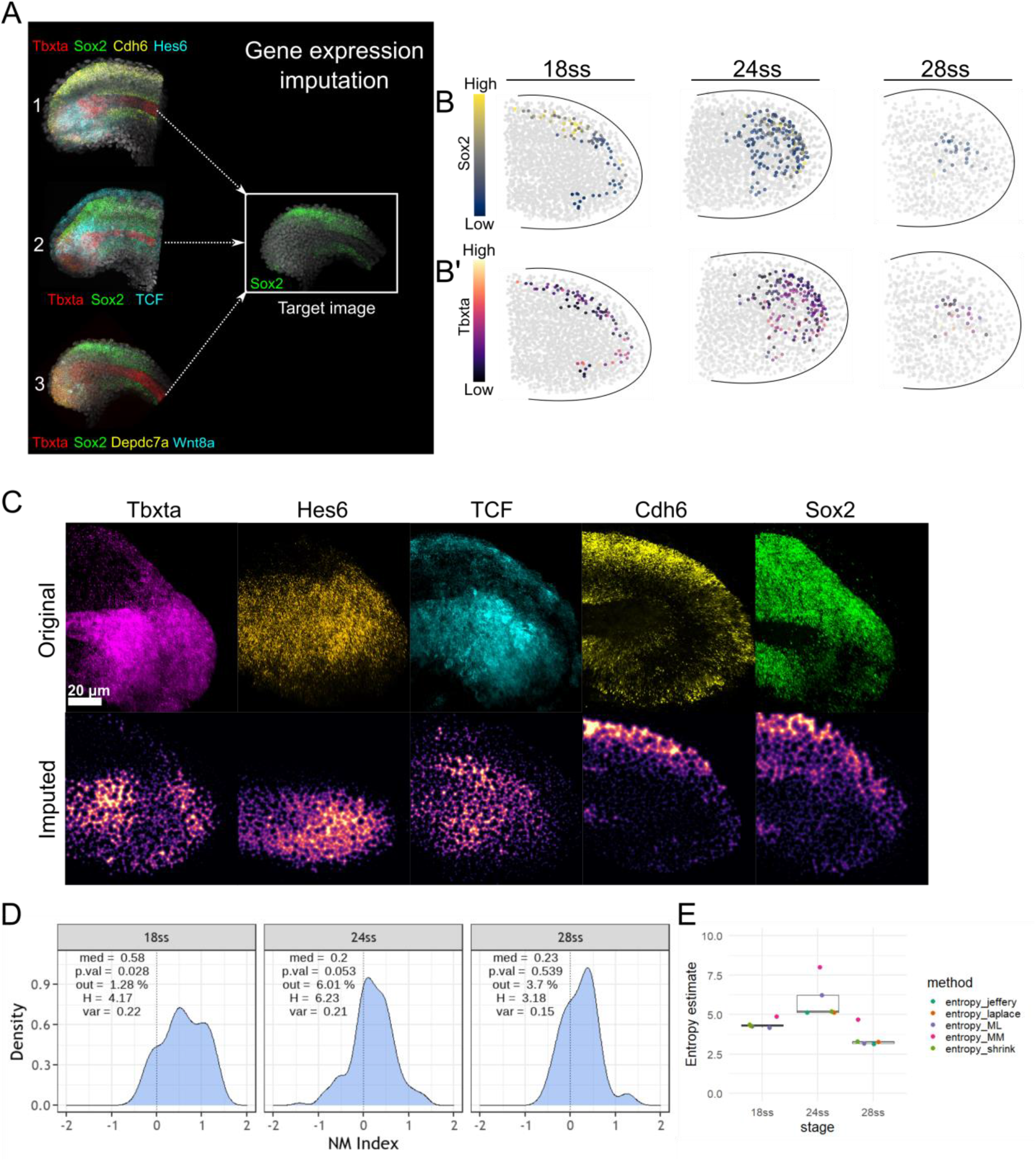
Gene expression imputation and the construction of a composite map via ZebReg demonstrates a peak in the NM index entropy at 24ss. (A) Application of ZebReg for the imputation of multiple genes onto a target composite image. In the panel, *tbxta, cdh6, hes6, tcf, depdc7a* and *wnt8a* are imputed onto a target image that is stained only for *sox2. sox2* is the common colour channel used to assist the alignment of the source images onto the target image. In this example, the resultant target image has 7 distinct colour channels. (B-B’) Colouring *in silico* NMps in the 18ss, 24ss and 28ss composite maps by (B) *sox2* expression (B’) *tbxta* expression levels. (C) The top ‘Original’ row depicts the 2D projections of the HCR data at 18ss. The bottom ‘Imputed’ row depicts the corresponding expression of these genes in the target composite image. (D) NM index density distributions computed from the 8-gene composite maps at 18ss, 24ss and 28ss. med: median; p.val: p-value for the Shapiro-Wilk test; out: outlier percentage; H: empirical entropy estimate; var: variance. (E) Entropy estimates of the NMp index, with the estimation of the standard error obtained via jackknife resampling. The entropy estimates consistently peak at 24ss. entropy_jeffrey: Dirichlet-multinomial pseudocount entropy estimator (Dirichlet) with Jeffrey’s prior; entropy_laplace: Dirichlet with Laplace’s prior; entropy_ML: empirical maximum likelihood estimator; entropy_MM: Miller-Madow estimator; entropy_shrink: James-Stein type shrinkage estimator.

We first used ZebReg to impute the expression of 8 genes into three composite maps, one for each stage at 18ss, 24ss and 28ss. Each composite map was constructed by combining the gene expression of 6 different images across different HCR experiments (Figure S8; Table S2), and each map displays the expression of *sox2* (Figure 3B), *tbxta* (Figure 3B’) and 6 other neural or mesodermal marker genes selected from our scRNA-seq analysis within a target point cloud. As these maps contain spatial information of the tailbud cells, we could identify the *in silico* NMps by virtue of their *sox2* and *tbxta* co-expression as well as their locations on the composite map (Figure S12). These *in silico* NMps in the composite maps were also found to be within the NMp regions in our probability map which were identified via a different approach (Figure S14).

To assess the fidelity of our gene expression imputation procedure, we performed a gene-by-gene qualitative inspection of the spatial expression patterns in the composite maps to the corresponding patterns observed in the original HCR images at 18ss. A visual comparison between the imputed and original images demonstrates a strong resemblance in their expression patterns (Figure 3C). For instance, *sox2* and *cdh6* are expressed strongly in the posterior neural tube and hypochord but not in the notochord. *tbxta* is expressed strongly in the notochord progenitor zone and the dorsal PW. Thus, the overall visual correspondence between the original and imputed gene expression images is evidence that ZebReg has aligned these images appropriately, at least when assessed on a qualitative level. We also performed additional *in silico* validation experiments and demonstrated that ZebReg also preserves the quantitative relationships between genes (See STAR Methods).

Using the composite maps, we first constructed an ‘NM index’ which combines the information across the 8 genes and plotted the NM index distributions for the NMps in the three composite maps (Figure 3D). These distributions reflect the neural/mesodermal biases of the *in silico* NMps in these three stages, and cells can be classified as being either neural-biased, mesoderm-biased or indecisive based on their NM index value (Figure S13A). We find that there is a consistent neural bias in the NMps across all three stages which is reflected by the median NM index value.

To quantify the NM heterogeneity of the *in silico* NMps between these stages, we computed a series of Shannon entropy estimators. Examining the empirical maximum-likelihood entropy estimator (H) with the natural unit of information (nat), we observed a surge in value at 24ss with H = 6.23 nat, compared to the neighboring values of H = 4.17 nat at 18ss and H = 3.18 nat at 28ss (Figure 3E). This increase in entropy followed by a decline was also observed in the other entropy estimates (Figure 3E). Thus, our data suggest that the NMp population heterogeneity, measured by the entropy, peaks at 24ss.

### Stochastic modelling of a genetic toggle switch with a time-varying input uncovers ‘rebellious’ cells

Our results indicate widespread cellular variability in the expression of NM genes during the process of NMp differentiation. Given the low-level expression of *sox2* and *tbxta* for the majority of NMps (Figure 1F), stochastic effects are likely to dominate and drive the cellular heterogeneity in the population (Kærn et al., 2005). To understand how noise-induced heterogeneities can arise in the context of the zebrafish tailbud, we first focused our analysis on constructing a single-cell resolution quantitative spatial landscape of the canonical Wnt signaling environment as it is one of the main signaling factors that bias the NM fate balance (Martin & Kimelman, 2012). We examined the nuclear expression of *tcf* mRNA (a readout of canonical Wnt) in the dorsal PW, intermediate zone and posterior PSM at the 18ss, 24ss and 30ss (Figure 4A-C). *tcf* expression was found to increase consistently along the dorsal PW to intermediate zone to PSM trajectory (Figure 4D), which traces the approximate path of the mesoderm-fated NMps.

**Figure 4.**
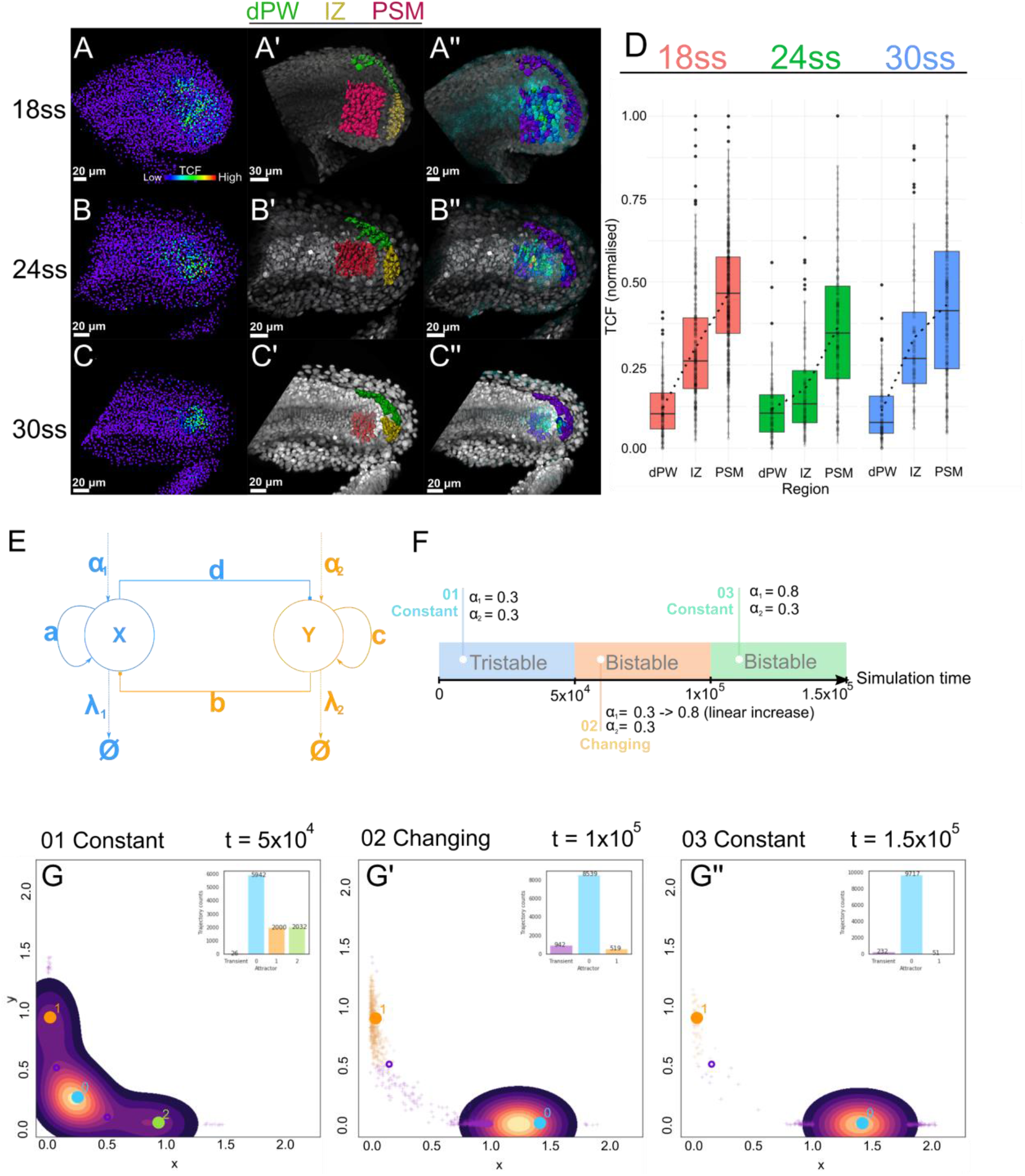
Stochastic modelling of a genetic toggle switch with a time-varying parameter uncovers ‘rebellious’ cells. (A-A’’) (A) Segmented 18ss tailbud nuclei coloured according to *tcf* expression. (A’) Nuclei residing in the dPW (green), IZ (yellow) and PSM (red) are depicted. (A’’) Same nuclei from (A’) coloured according to *tcf* expression. dPW: dorsal posterior wall, IZ: intermediate zone, PSM: presomitic mesoderm. (B-B’’) Similar to (A-A’’) showing tailbuds at 24ss. (C-C’’) Similar to (A-A’’) showing tailbuds at 30ss. (D) Quantification of normalised *tcf* expression levels in the dPW, IZ and PSM at 18ss, 24ss and 30ss. (E) Schematic representation of the toggle switch model. (F) Schematic of the computational experiment. The experiment has three stages: first (01 constant), the simulations are allowed to equilibrate in the tristable dynamical regime from t = 0 to t = 5×10^4^. Next (02 changing), from t = 5×10^4^ to t = 1×10^5^, the external activation parameter, α_1_ is increased stepwise and reaches a value of 0.8 at t = 1×10^5^. Finally (03 constant), α_1_ is kept constant for the remaining simulation duration from t = 1×10^5^ to t = 1.5×10^5^. (G-G’’) Combined results of 10,000 stochastic simulations. Each ‘+’ symbol corresponds either to the location of a ‘transient’ trajectory with transient behaviour (purple) or to a ‘rebellious’ trajectory residing in the alternative attractor (orange). By t = 1.5×10^5^ (end of 03 Constant), most trajectories have entered the primary attractor (attractor 0).

To incorporate the changes in Wnt signalling levels as NMps differentiate into their mesodermal fates, we constructed a stochastic non-autonomous genetic toggle switch model of *sox2* and *tbxta*, where both genes mutually repress each other (Figure 4E). Whilst a direct repression between *tbxta* and *sox2* has not been shown, the interaction between *sox2* and *tbxta* in other NMp systems, whether direct or indirect, is an antagonistic one that drives the mutually exclusive choice between the neural or mesodermal fate (Bouldin et al., 2015; Gouti et al., 2017).

When stochastic trajectories were first equilibrated within a tristable regime in the first interval of our simulation (Figure 4F, 01 Constant), they spread out to occupy all three attractors with considerable frequency (Figure 4G). To reflect the increase in Wnt signalling within the differentiating NMps along the mesodermal trajectory, we then increased the external activation parameter α_1_ linearly over the next simulation interval (Figure 4F, 02 Changing). At the end of the second interval, we observed that most stochastic trajectories reside within the primary ‘mesodermal’ attractor (attractor 0) (Figure 4G’). However, some trajectories deviate from this behaviour - around 9.5% of trajectories exhibit transient behaviour and are situated between both neural and mesodermal attractors. Unexpectedly, a smaller percentage of trajectories (5.1%) still reside within the alternate ‘neural’ attractor (attractor 1). In the final simulation interval (03 Constant) during which the external activation parameter α_1_ is kept constant at a value of 0.8 (Figure 4G’’), these trajectories eventually switch into the primary attractor and adopt the appropriate cell fate.

Thus, the introduction of non-autonomy into a stochastic toggle switch leads to a novel phenomenon that was not present in the autonomous case (Figure S16B) - the emergence of ‘rebellious’ cells with gene expression states (high *sox2*, low *tbxta*) inconsistent with their signalling environment (high Wnt) and eventual mesodermal fate.

### ZebReg’s composite maps reveal that the number of Rebellious cells peak at 24ss

We returned to ZebReg’s composite maps to assess whether these Rebellious cells can be found within the *in silico* NMp population. To identify these cells, we need to compare their canonical Wnt signalling activities (*tcf* levels) and eventual cell fates (neural or mesodermal) against their NM gene expression states (NM index levels). Given that our data consist of fixed snapshot images of the NMps, we do not have direct information of their prospective fates. Despite this, as NMps at the 21ss onwards are spatially segregated and have low levels of proliferation (Attardi et al., 2018, Figure S11A-B, D-E), we reason by an argument of elimination that since NMps also display low levels of apoptosis throughout 18ss to 30ss (Figure S11A,C), we can reliably predict the cell fate of NMps based on their spatial locations in the composite maps at 18ss, 24ss and 30ss.

Having established that we can infer an *in silico* NMp’s fate prospectively from its location in the composite map, we defined approximate neural-fated and mesodermal-fated domains following the fate map of Attardi and colleagues and assessed the NM index levels of the NMps within these domains (Figure 5A). We found cells with NM gene expression profiles that are inconsistent with their prospective fates and labelled these cells as ‘Incongruent’. Conversely, cells with compatible state-fate relationships are labelled ‘Congruent’. We observed Incongruent cells with low/high NM index levels residing in the neural/mesoderm-fated domains (Figure 5A blue arrows). As *sox2* and *tbxta* are the primary orchestrators of the neural and mesodermal gene expression programmes respectively, these ‘incongruent’ cells are also found in our original HCR images as *sox2* (*tbxta*)-high NMps within the neural (mesoderm)-fated domains (Figure 5B).

**Figure 5.**
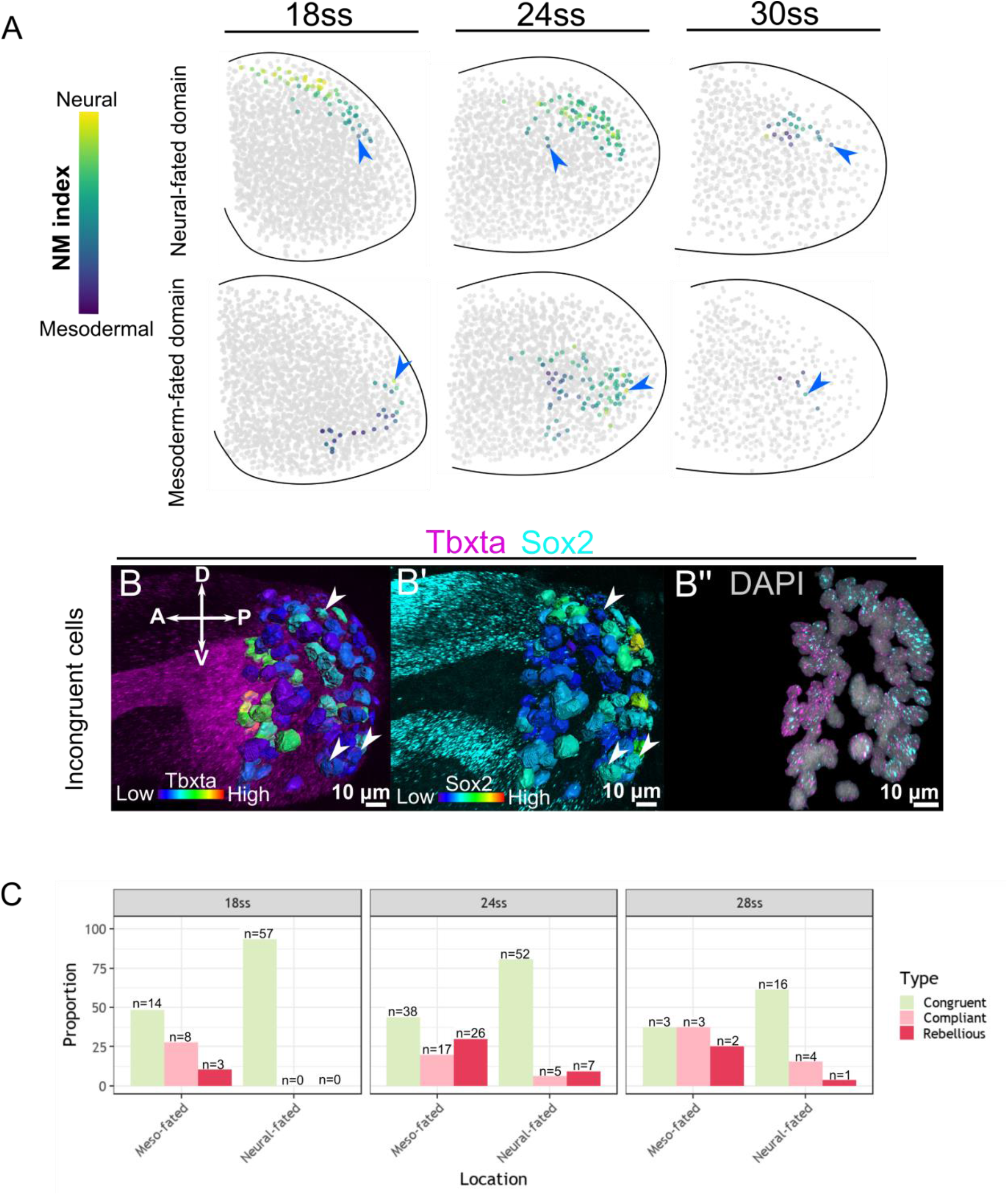
ZebReg’s composite maps reveal that the number of Rebellious cells peak at 24ss. (A) Demarcation of the neural-fated and mesoderm-fated domains in the composite maps. Non-NMps are coloured grey, whereas NMps are coloured according to their NM index levels. Blue arrows mark incongruent cells. (B-B’’) HCR stains of a representative zebrafish tailbud at 24ss for *tbxta* and *sox2*. Segmented surfaces correspond to NMps which are coloured by the expression levels of *tbxta* (B) and *sox2* (B’). Nuclear signals for *sox2* and *tbxta* are shown to illustrate co-expression (B’’). Arrow heads mark Incongruent cells. (C) Proportion of Congruent, Compliant and Rebellious cells in the mesoderm-fated and neural-fated domains at 18ss, 24ss and 28ss. At each stage, summing up the number of Compliant and Rebellious cells yields the number of Incongruent cells.

Incongruent cells can be further classified as ‘Compliant’ or ‘Rebellious’ depending on whether their Wnt signalling activities (*tcf* expression levels) are consistent or inconsistent with their NM gene expression states (Table S1). We quantified the proportion of Compliant, Rebellious and Congruent cells in the mesodermal and neural-fated domains of our three composite maps (Figure 5C). At 18ss and 28ss, most cells are Congruent. Also, more Incongruent cells are found in the mesoderm-fated domain than the neural-fated domain. However, at the 24ss, we find a greater number of Incongruent cells (Compliant and Rebellious) than Congruent cells in the mesoderm-fated domain. Specifically, the number of Rebellious cells in the mesoderm-fated domain peaks at this stage. Thus, consistent with the transition state model, we find a loosening of the relationship between cell state and fate as reflected by the increase in the number of rebellious NMps at the 24ss prior to their commitment to the NM fate.

## Discussion

Zebrafish tailbud NMps have proven to be an attractive *in vivo* system to assess the transition state hypothesis. Specifically, we investigated whether a transient window of elevated stochasticity in gene expression precedes the NMp differentiation event at around 24ss.

Our discovery of rebellious cells in the ZebReg composite maps recapitulates the finding by Mojtahedi and colleagues in their *in vitro* study on the differentiation of a multipotent hematopoietic cell line (Mojtahedi et al., 2016). Rebellious cells emerge at day 3 post-treatment as cells that express an erythroid profile when stimulated with Granulocyte macrophage colony-stimulating factor /IL-3 or a myeloid profile when stimulated with erythropoietin. Eventually, cells disappear at day 6 post-treatment. In addition, ‘edge’ cells were identified in cancer cell lines as cells that adopt a gene expression profile that is different from the average profile in the population distribution (Li et al., 2016). This phenomenon is not unprecedented *in vivo*. In an earlier study on Xenopus embryonic development (Wardle & Smith, 2004), cells that express a lineage marker at the ‘wrong’ place, such as Goosecoid expressing cells in the ventral instead of the dorsal region of the embryo, were labelled as ‘rogue’ cells to indicate their abnormal expression profile. These cells appear more frequently in the early gastrula stage and reduce in frequency at the late gastrula stage. In both cases, these rebellious/rogue cells are proposed to ‘fit in or die trying’ - they would either die by apoptosis or transdifferentiate to adopt the appropriate gene expression profile if rescued by delivery of the appropriate signal or through interactions with neighbouring cells via the community effect.

In the context of zebrafish NMp differentiation, our finding that the proportion of rebellious cells (Figure 5C) is highest at 24ss within the mesoderm-fated domain extends a recent work on the connection between morphogenetic movements and mesoderm fate acquisition in the zebrafish NMps (Kinney et al., 2020). *Sox2* and canonical Wnt co-expression in mesoderm-fated NMps primes these cells towards both neural and mesodermal fates and acts as a developmental checkpoint that traps these cells in a poised, intermediate state. This intermediate state where EMT is delayed resembles a hybrid EMT transition state found in cells with high potential for metastasis (Yang et al., 2020). In fact, *tbxta* (Brachyury) is a driver of EMT in various tumors and is correlated with metastatic activity and the acquisition of a mesenchymal phenotype (Chen et al., 2020). Thus, our work emphasises a strong connection between the hybrid EMT transition state expressing multiple intermediate cell states (Sha et al., 2019) and the neural-mesodermal transition state (Steventon & Martinez-Arias, 2017). The correspondence between morphological fluctuations and the entry into a transitory state was also recently proposed in a study on hematopoietic stem and progenitor cells (Moussy et al., 2017). As NMps exit the transition state and differentiate into the neural or mesodermal fates, the heterogeneity in *sox2* and *tbxta* expression is resolved as NMps adopt either a high *sox2*/low *tbxta* (neural) or high *tbxta*/low *sox2* (mesodermal) expression profile (Figure 1D-F). This is consistent with the proposed role of Sox2 and Bra protein level ratios dictating the specific cell movements associated with each lineage (Romanos et al., 2021).

Recent work on multipotent zebrafish neural crest cells suggests that at least a portion of the neural biased trunk neural crest (NC) progenitors arise from early neural biased zebrafish NMps at 5-6ss (Lukoseviciute et al., 2021). A similar conclusion was reached in multiple studies of in vitro human pluripotent stem cell-derived axial progenitors, demonstrating that the generation of trunk NCs involves an obligatory NMp intermediate (Frith et al., 2018; Hackland et al., 2019). We identified *sp5l, cdh6, znf703* and *foxd3* as differentially expressed genes of the NMp cluster; all of which have important roles in neural crest specification. In addition, many differentially expressed genes identified from the NMp cluster are involved in signalling pathways (FGF, Wnt, BMP) and the synergistic action of these pathways play a critical role in neural crest differentiation (Sauka-Spengler & Bronner-Fraser, 2008). When we photolabelled the dorsal PW (NMp region) at 18ss and tracked these cells until 28ss, we noticed that the anterior photolabels in the dorsal neural tube appear to be emigrating away whilst the posterior labels do not show signs of migration (Figure S11D). 24 hours later, the photolabels were found to have spread more anteriorly, with the anterior labels appearing more dispersed ventrally. When we photolabelled the dorsal PW at 28ss, we also found similar photolabels in the dorsal neural tube 24 hours post-photolabeling (Figure S11E). The localisation of the labels in the dorsal neural tube alongside the anterior pattern of cell migration strongly suggest that the differentiation of the NMp-derived neural progenitors into the trunk NC progenitors continues throughout somitogenesis and occurs even as we approach the end of somitogenesis. Therefore, we extend the observation made by Lukoseviciute and colleagues, providing support for an NMp to trunk NC progenitor lineage that occurs even in the later tailbud NMp population.

To the best of our knowledge, our work is the first to directly catalog the transient surge in heterogeneity in mRNA expression *in vivo* during an endogenous differentiation event in a wild-type vertebrate species. Whilst several studies have proposed mechanistic models to explain the relationship between transcriptional heterogeneity and cell fate commitment (Antolović et al., 2017; Pina et al., 2012) and even functional pluripotency (MacArthur & Lemischka, 2013), our study was not designed to discriminate between these causal models. Instead, we focused on assessing the association between cell fate commitment and the increase in gene expression heterogeneity *in vivo*. Future work, outside the scope of this paper, is necessary to fill in the mechanistic details that generate these heterogeneities during cell fate transitions *in vivo*.

Taken together, our work supports the existence of a transition state within an endogenous cell fate decision making event. Recognising the functional importance of transcriptional stochasticity and non-genetic heterogeneities during differentiation has important practical consequences. It drove the discovery that regulators of transcriptional noise may play a general role in the acquisition of malignancy by modulating the balance between proliferation and differentiation (Domingues et al., 2020), and may be an important dimension to consider when improving the efficacy of mesenchymal stem cell-based therapies (McLeod & Mauck, 2017; Pacini, 2014). Seen alongside the evidence presented from other systems, it becomes increasingly plausible that the transition state is not an idiosyncrasy of *in vitro* culture conditions or a peculiarity of cancer models. We await future developments on whether the critical behaviours predicted in the transition state model are a universal characteristic of cell state transitions in biological systems *in vivo*.

## Acknowledgements

We would like to thank Nick Monk for advice on the computational modelling project. We also thank all members of the Steventon lab for comments on the manuscript. K.T. is funded by the Cambridge Commonwealth, European & International Trust under a Cambridge International Scholarship. D.S. was supported by the Wellcome Trust-funded Developmental Mechanisms PhD programme (220022/Z/19/Z). B.V. was supported by the Department of Zoology at the University of Oxford. B.J.S. was supported by a Henry Dale Fellowship jointly funded by the Wellcome Trust and the Royal Society (109408/Z/15/Z).

## Author contributions

K.T. and B.S. designed the study and wrote the manuscript; K.T. and D.S. jointly developed ZebReg; K.T. performed the experiments and analysed the data; B.V. provided advice on the computational modelling approach. All authors read and approved the final manuscript.

## Limitations of the study

The coloured ICP (cICP) algorithm employed in ZebReg will not be able to align point clouds exactly, as zebrafish tailbuds will inevitably differ from one another in their nuclei position and gene expression intensities. Instead, for each nucleus from the source image, ZebReg can, at best, map it to its most similar cell counterpart in the target image, based on their proximity to each other and similarity in expression of a reference gene. For multiply mapped and unmapped target points, ZebReg imputes their gene expression intensities by taking the average intensities of each point’s k-nearest neighbors (k=5). This approach assumes a degree of spatial autocorrelation in gene expression intensities of a point with its neighbours. During the transition state where cell-cell correlation decreases, our approach may underestimate the extent of cellular heterogeneity in the population due to the application of an averaging procedure.

In our work, we adopted a descriptive, fixed imaging-based approach towards interrogating the level of gene expression heterogeneities during NMp differentiation. Whilst the peculiarities of the zebrafish NMp model enabled us to infer cellular fates from cellular positions, without adopting a live imaging approach, we were unable to document the details of the transcriptional dynamics around 24ss (Weinreb et al., 2017). Zebrafish embryos have been amenable to live RNA imaging using various techniques such as the MS2 labelling system (Campbell et al., 2015), 3’ poly(A) tail labelling system (Westerich et al., 2020) and molecular beacon sensors (Li et al., 2017) due to its optical transparency. Thus, future work could perform live imaging of *sox2* and *tbxta* mRNAs to monitor the changes in transcription dynamics around 24ss as NMps enter and exit the transition state.

## STAR Methods

### Key Resources Table

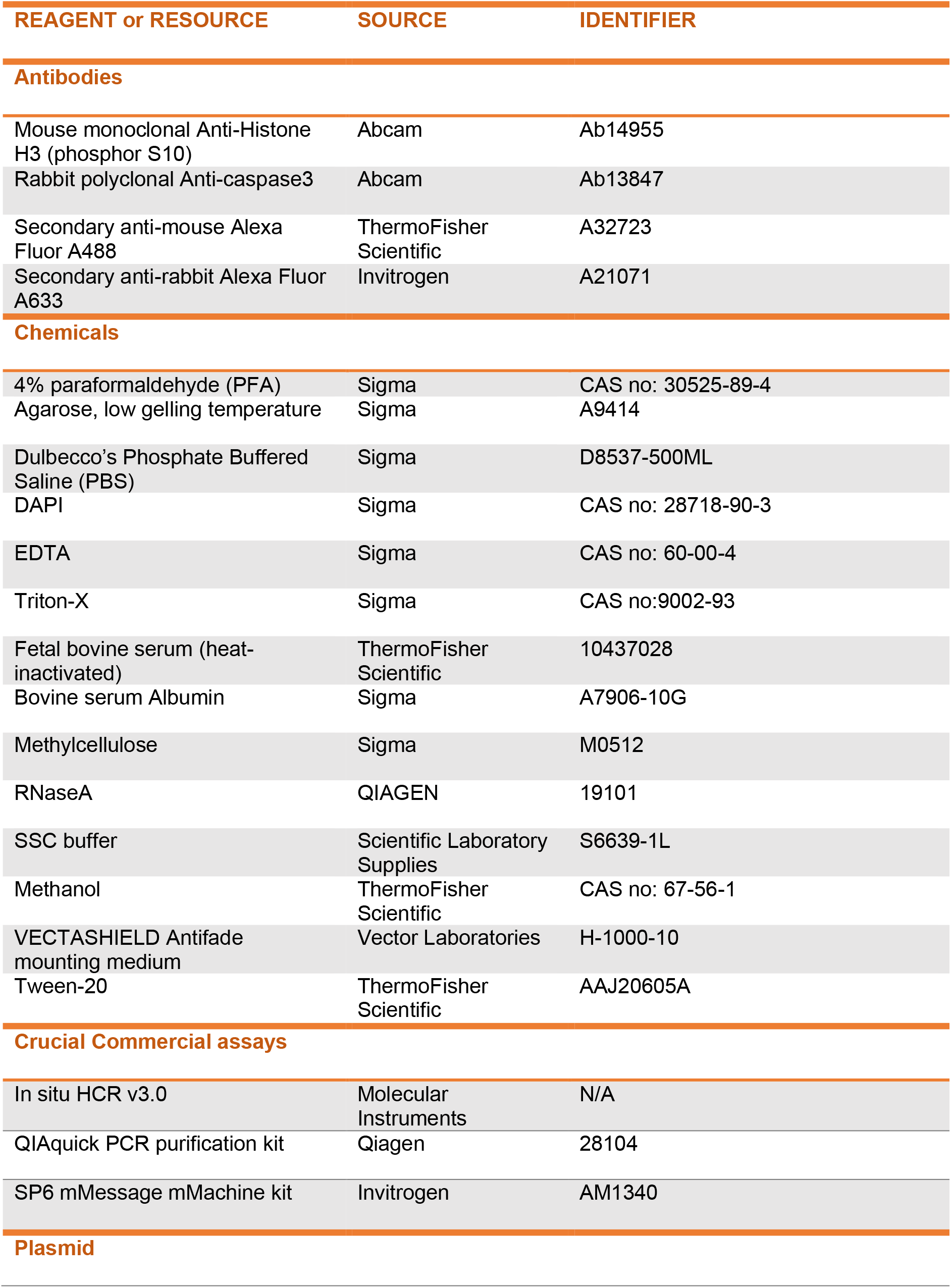
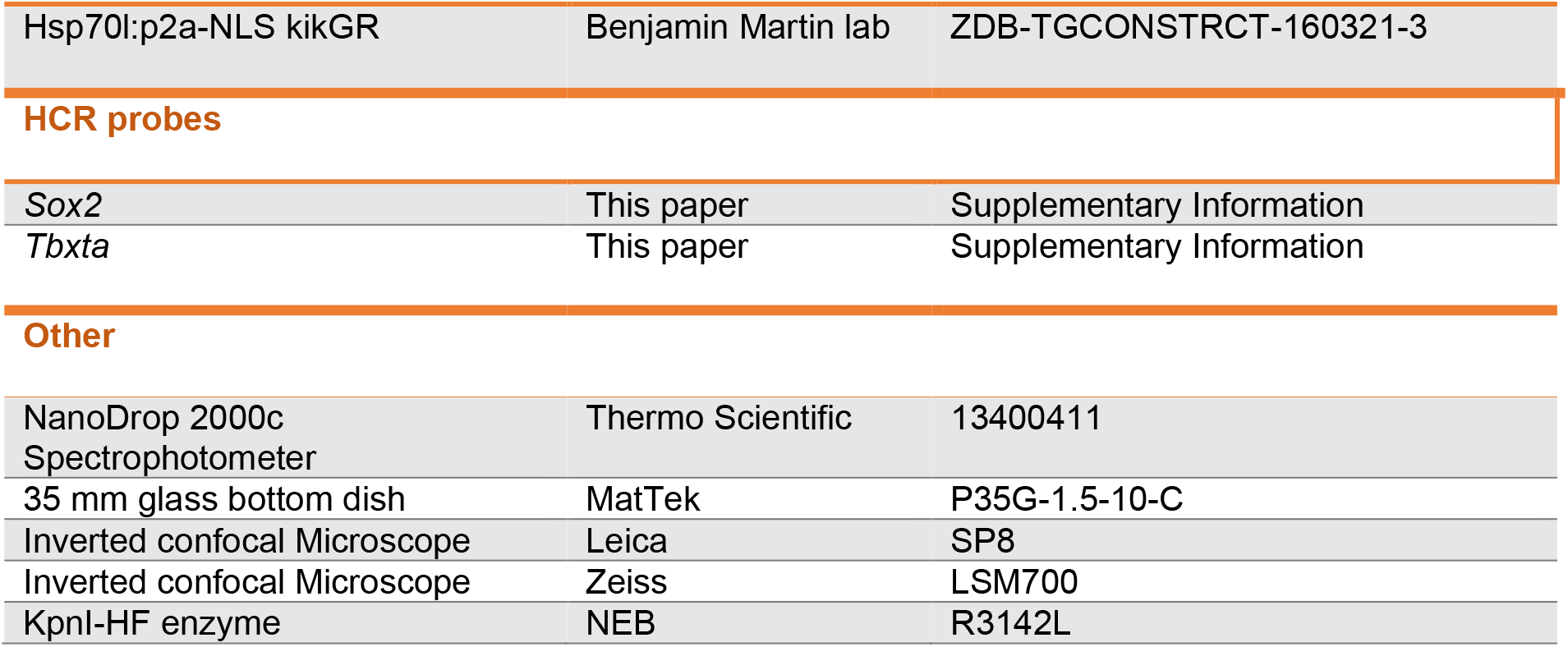

| REAGENT or RESOURCE | SOURCE | IDENTIFIER |
| --- | --- | --- |
| **Antibodies** |  |  |
| Mouse monoclonal Anti-Histone H3 (phosphor S10) | Abcam | Ab14955 |
| Rabbit polyclonal Anti-caspase3 | Abcam | Ab13847 |
| Secondary anti-mouse Alexa Fluor A488 | ThermoFisher Scientific | A32723 |
| Secondary anti-rabbit Alexa Fluor A633 | Invitrogen | A21071 |
| **Chemicals** |  |  |
| 4% paraformaldehyde (PFA) | Sigma | CAS no: 30525-89-4 |
| Agarose, low gelling temperature | Sigma | A9414 |
| Dulbecco's Phosphate Buffered Saline (PBS) | Sigma | D8537-500ML |
| DAPI | Sigma | CAS no: 28718-90-3 |
| EDTA | Sigma | CAS no: 60-00-4 |
| Triton-X | Sigma | CAS no:9002-93 |
| Fetal bovine serum (heat-inactivated) | ThermoFisher Scientific | 10437028 |
| Bovine serum Albumin | Sigma | A7906-10G |
| Methylcellulose | Sigma | M0512 |
| RNaseA | QIAGEN | 19101 |
| SSC buffer | Scientific Laboratory Supplies | S6639-1L |
| Methanol | ThermoFisher Scientific | CAS no: 67-56-1 |
| VECTASHIELD Antifade mounting medium | Vector Laboratories | H-1000-10 |
| Tween-20 | ThermoFisher Scientific | AAJ20605A |
| **Crucial Commercial assays** |  |  |
| In situ HCR v3.0 | Molecular Instruments | N/A |
| QIAquick PCR purification kit | Qiagen | 28104 |
| SP6 mMessage mMachine kit | Invitrogen | AM1340 |
| **Plasmid** |  |  |
| Hsp70l:p2a-NLS kikGR | Benjamin Martin lab | ZDB-TGCONSTRCT-160321-3 |
| **HCR probes** |  |  |
| *Sox2* | This paper | Supplementary Information |
| *Tbxta* | This paper | Supplementary Information |
| **Other** |  |  |
| NanoDrop 2000c Spectrophotometer | Thermo Scientific | 13400411 |
| 35 mm glass bottom dish | MatTek | P35G-1.5-10-C |
| Inverted confocal Microscope | Leica | SP8 |
| Inverted confocal Microscope | Zeiss | LSM700 |
| KpnI-HF enzyme | NEB | R3142L |

### Resource Availability

#### Lead contact

Further information and requests for resources should be directed to and will be fulfilled by the lead contact, Ben Steventon.

#### Materials availability

HCR probe sequences for *sox2* and *tbxta* are documented in the supplementary file.

#### Data and code availability

- Code for establishing ZebReg can be found here: https://github.com/DillanSaunders/ZebReg
- Any additional information required to reanalyze the data reported in this paper is available from the lead contact upon request.

### Experiment Model and Subject Details

#### Zebrafish husbandry

All zebrafish procedures were conducted under the Animals (Scientific Procedures) Act 1986 Amendment Regulations 2012, following ethical review by the University of Cambridge Animal Welfare and Ethical Review Body (AWERB). Wild type lines used are either Tüpfel long fin (TL), AB/TL or AB. The Tg(7x*TCF*-Xla.Sia:GFP) reporter line (Moro et al., 2012) was provided by the Steven Wilson laboratory. All embryos obtained were obtained and raised in standard E3 media at 28ºoC. Embryos were staged according to Kimmel et al., 1995.

### Method Details

#### Version 3 Hybridisation Chain Reaction (V3 HCR)

Zebrafish embryos at the required stages were fixed in 4% PFA in DEPC-treated, calcium and magnesium-free PBS at 4 ^°^ C overnight. Embryos were then stained with V3 HCR (Choi et al., 2018). All hairpins were purchased from Molecular Instruments. All probes were purchased from Molecular Instruments except for *sox2* and *tbxta* which were manually designed. After the staining procedure, samples were counterstained with DAPI at a dilution of 1:1000 in 5xSSCT for 2 hours at room temperature. The tailbud region was cut out with a forceps and eyelash tool, and then mounted on a 35mm glass bottom dish (MatTek) with the Vectashield antifade mounting medium for confocal imaging.

#### Quantification of nuclear gene expression intensities in NMps

HCR images were processed in Imaris (Bitplane). Unless otherwise stated, all *sox2*+*tbxta*+ HCR images were analysed for the number of NMps as described in Figure S1.

#### Immunostaining with anti-PH3 and anti-caspase 3 antibodies

Zebrafish embryos at the required stages were fixed in 4% PFA in DEPC-treated, calcium and magnesium-free PBS at 4oC overnight. Embryos were then co-stained with a 1:500 dilution of mouse anti-PH3 antibody (abcam, ab14955) and 1:500 dilution of rabbit anti-caspase3 antibody (abcam, ab13847), as described in Sorrells et al., 2013. Secondary anti-mouse Alexa Fluor 488-conjugated antibody and anti-rabbit Alexa Fluor 647-conjugated antibody were both diluted in 1:500 PDT solution and incubated with the samples overnight at 4°C. DAPI was added at the final step with a 1:1000 dilution in PDT and incubated for 2 hours at room temperature for nuclear detection. Images were quantified in the 3/4D Image Visualisation and Analysis Software Imaris 9.2.1 (Bitplane). The percentages of mitotic or apoptotic cells for each sample were calculated as the fraction of PH3+ or caspase3+ nuclei over the total number of nuclei in the tailbud, multiplied by 100.

#### Photolabeling with nuclear-targeted kikume

The hsp70l:p2a-NLS kikGR vector (Bouldin et al., 2015) was extracted from an overnight grown bacterial culture. Briefly, bacterial cells were collected via centrifugation and washed sequentially with the following 3 buffers: P1 containing 50 mM Tris-Cl at pH 8.2, 10 mM EDTA at pH 8.0, RNase A (QIAGEN); P2 (filter-sterilised) containing 0.8% NaOH and 1% SDS; P3 containing 3M KOAc that is adjusted to pH 5.5 with glacial acetic acid. Plasmid DNA was precipitated with 70% isopropanol and washed with 70% ethanol before resuspension in nuclease-free water.

The vector was linearised by restriction digestion with the KpnI-HF enzyme (NEB), and subsequently purified using the QIAquick PCR purification kit (Qiagen). The purified, linearised plasmid was transcribed at the SP6 promoter with the SP6 mMessage mMachine kit (Invitrogen), and lithium chloride precipitation was carried out for mRNA recovery. Quantification of the transcribed kikGR mRNA was performed on the NanoDrop instrument (Thermofisher).

One-cell stage zebrafish embryos were injected with the NLS-kikGR mRNA and then embedded in low gelling point agarose (Sigma) at 1% w/v in E3 media at the bottom of a MatTek 35mm glass bottom dish. Photoconversion and image acquisition was performed on a Zeiss LSM 700 confocal microscope. Efficient, irreversible photoconversion of NLS-KikGR in the zebrafish embryos at mid-somitogenesis stages was carried out by scanning the 405 nm laser at 15% laser power for approximately 30 seconds in a region of interest.

#### Confocal microscopy imaging

Samples were imaged on either a Zeiss LSM700 inverted confocal or a Leica TCS SP8 inverted confocal at 10X, 20X or 40X magnification.

#### Analysis of scRNA-seq data

##### Downloading the 18hpf scRNA-seq dataset and preprocessing

The wild type 18hpf zebrafish scRNA-seq raw counts dataset and the associated clusterIDs were downloaded from GEO with the accession number GSM3067194 (Wagner et al., 2018). First, outlier cells with log-transformed library and feature sizes more than 3 median absolute deviations (MADs) from the respective median metric values were removed. Genes that were not expressed in the dataset were filtered out. At this quality control threshold, most genes and cells were retained for downstream analysis, resulting in a dataset with 30296 genes x 6954 cells (381 genes and 8 cells discarded). The data was then converted into a Seurat 3.0 object (Stuart et al., 2019) for subsequent analyses. Cell cycle scoring and regression were performed in Seurat 3.0 using a set of cell-cycle associated genes for zebrafish (Lush et al., 2019), with the S.Score and G2M.Score as inputs to the vars.to.regress argument in the *SCTransform* function. Data normalisation, scaling and the identification of the top 3000 most variable genes were also carried out using the *SCTransform* wrapper.

##### Low dimensional embedding and Louvain clustering

The normalised and scaled data was projected into low dimensional subspace via principal components analysis (PCA) with default settings for the *RunPCA* function. (Figure 2.2B). Following this, the uniform manifold approximation and projection (UMAP) embedding was implemented via the *RunUMAP* function. To perform clustering, groups of similar cells on the UMAP embedding were identified by generating a shared nearest neighbor (SNN) graph of the dataset with the *FindNeighbors* function, and then clustered using the Louvain algorithm with the *FindClusters* function at various resolutions. Subclustering on the tailbud cells was performed in similar manner to the above clustering procedure, with a resolution of 1 set for the FindClusters function. To examine the clustering results, clustering trees were plotted with the *clustree* package whilst the adjusted rand index and clustering entropy were implemented in the *mclust* and *NMF* packages respectively.

##### Identification of differentially expressed genes

For each cluster, supervised annotation was carried out by examining the marker genes identified by a Model-based Analysis of Single-cell Transcriptomics (MAST) and a Wilcoxon Rank-Sum test. The tests were carried out using the *FindAllMarkers* function in Seurat that compares cells in each cluster against all other remaining clusters. The function is set to return only positive markers for each cluster (only.pos = TRUE). Differentially expressed genes with an adjusted p-value less than 0.05 were retained for analysis. They were then sorted in order of priority, based on the log fold-change of the average expression between the cluster under study and the remaining 7 tailbud subclusters (avg_log2FC).

##### Robustness analysis of tailbud clustering assignments

To assess the robustness of our selection of the zebrafish tailbud cells from the 18hpf dataset, we employed a different approach than Wagner and colleagues (Wagner et al., 2018) by embedding the 6,954 cells in the 18hpf dataset into a Uniform Manifold Approximation and Projection (UMAP) space and using the Louvain community detection algorithm to identify clusters (Figure S3A).

We first assessed the similarity between the two data clusterings using the Adjusted Rand index (ARI) and clustering entropy index. High ARI values and low entropy values are obtained across a wide range of clustering resolutions, apart from the initial resolution of 0.2 (Figure S4A,C). In addition, analysis of the clustering tree shows that at a resolution of 0.2, there are 11 clusters which continue to be split up gradually. At increasing resolutions, the number of in-proportion edges (edges with low transparency) remain low which indicates only minor changes in the clustering tree. At a clustering resolution of 0.8, we obtained 22 clusters (Figure S4B). When we re-examined the distribution of our tailbud labels against Wagner and colleagues’ labels, we find that they are highly concordant (Figure S3B), suggesting that our selection of the zebrafish tailbud cells are robust across different analytical strategies. As the Louvain algorithm is stochastic, we re-ran the algorithm for 10 iterations and retained cells that are consistently located in the tailbud clusters for 9 and 10 iterations for downstream analyses (Figure S4D).

##### Computation of the critical index and transcriptional noise index

The critical index is defined as the ratio of two averaged Pearson correlation coefficients: the average correlations between all pairs of gene vectors over the average correlations between all pairs of cell state vectors (Mojtahedi et al., 2016). In the scRNA-seq analysis, to account for the differences in cell number between clusters, 200 cells from each cluster were randomly sampled with replacement to calculate the index, and the procedure was repeated for 10,000 times. We also assessed the robustness of the critical index to differences in cell number and number of marker genes used (Figure S5).

The transcriptional noise index was measured using the top 2000 highly variable genes of each cluster following the work of Mohammed and colleagues (Mohammed et al., 2017).

#### Tailbud image registration with ZebReg

##### Overview of pipeline

ZebReg is a 3D, non-landmark-based image registration Python tool which we developed to integrate cellular position and nuclear gene-expression information from confocal images of zebrafish tailbuds. Leveraging upon the open-source open3D library (Zhou et al., 2018), ZebReg implements a set of rigid body, point-based registration algorithms that are popular in the field of geometric registration to align a 3D point cloud (source cloud) into a reference point cloud (target cloud).

At present, we have tested ZebReg on zebrafish tailbuds ranging from 18ss to 30ss. Briefly, confocal images were first preprocessed in Imaris to obtain segmented DAPI-stained nuclei (Figure S1). Next, to ensure a consistent field of view, all nuclei posterior to the tip of the developing notochord for all the images were retained for analysis. ZebReg performs the alignment by first importing the 3D centroid coordinates and gene expression intensities (if present) of the segmented nuclei and converting each image into a point cloud. Then, given a set of source clouds and a reference point cloud (target cloud), ZebReg finds the best linear transformation (no shearing, stretching or other deformations) between each source cloud and the target cloud. Additionally, if colour intensities of the source clouds are provided, ZebReg can map them onto the target cloud by imputing the gene expression intensities in the target point cloud and thus generate a composite image (See Figure S7)

##### Imputation of gene expression intensities

There are three possible sets of outcomes during the imputation procedure:

i. First, the mapping of the source point to the target point may be unique, in which case the target point simply adopts the intensity value of the corresponding source point.
ii. In cases where there is not a single source point corresponding to the target point, ZebReg provides the user with several options to resolve the discrepancy. If multiple source points map to the same target point, the target point adopts either the mean or median of these source intensity values (default: ‘median’).
iii. Alternatively, if there is no source point that corresponds to the target point, ZebReg provides three options to impute the gene expression intensity of this target point: ‘null’, ‘complete’ or ‘knn’ (default : ‘knn’). ‘null’ sets the intensity of the target point to 0, whilst ‘complete’ can be used if the target point cloud already has an intensity channel for that gene, in which case the point simply retains the original target intensity value. In the default ‘knn’ case, regression is performed based on the k-nearest neighbors of the point (default : n = 5) as implemented in the *sklearn* package. The target point takes on the mean intensity value of the closest target points.

Notably, in cases ii) and iii), ZebReg imputes the expression intensities of the target points by borrowing information from multiple source or neighboring target points.

##### Point set registration algorithms

To conduct the image alignment, ZebReg employs the following point set registration algorithms:

i. Random Sample Consensus (RANSAC) The RANSAC algorithm is a non-deterministic global alignment algorithm that is used in ZebReg to provide the initial coarse alignment for the ICP and cICP local algorithms (Fischler & Bolles, 1981).
ii. Iterative Closest Point (ICP) In the vanilla ICP algorithm, the algorithm repeatedly updates the transformation required to map the source to target cloud by minimizing the distance between points (Besl & McKay, 1992). In ZebReg, we use the point-to-plane ICP variant due to its increased speed of convergence (Rusinkiewicz & Levoy, 2001).
iii. Coloured Iterative Closest Point (cICP) For images with a colour channel in common, it is advantageous to consider their colour on top of geometry during point set registration. In these cases, ZebReg uses cICP, a modified version of ICP implemented in open3D, which optimises a joint geometric and photometric objective (Park et al., 2017).

ZebReg carries out all alignments by first performing a coarse global alignment with RANSAC, followed by either a finer alignment with ICP (if no colour channel is supplied) or cICP (if a common colour channel is present in the source and target image).

##### In silico validation of ZebReg

First, we constructed a mean absolute error (MAE) metric which quantifies the average difference in normalised signal intensities of the shared colour channel between the source and target image pair after image registration. To assess the accuracy of ZebReg’s image alignment with the cICP algorithm, we selected a source and target point cloud of the zebrafish tailbud and used the MAE as the test statistic in a permutation test which tests the following hypotheses:

H_0_ : ZebReg cICP registration has no effect on the colour intensity residuals between source and target clouds.

H_1_ : ZebReg cICP registration reduces the colour intensity residuals between source and target clouds.

In the permutation test, the sampling distribution under the null hypothesis was constructed by randomly rearranging the order of the target colour array, and then calculating the MAE using the permutated and original target colour arrays over 10,000 iterations. In effect, the null distribution provides the range of MAE estimates under the condition where ZebReg’s reported correspondence mapping between the source and target colour arrays is random. The null distribution was then fit to a gaussian distribution for the computation of the 95% confidence interval (Figure S9A).

Next, to assess the effectiveness of the various point cloud registration algorithms, we registered a point cloud with its rotated counterpart using 3 algorithms that are implemented in open3D: RANSAC, ICP and cICP, and assessed whether they can successfully recover the correspondence map. (Figure S9B)

We then compared the cICP’s performance across the different datasets. The datasets we have chosen for comparison were (Figure S9C-D):

- *Test sample*: Images of two separate zebrafish tailbuds at 18ss. Test sample exemplifies the performance of the algorithm on an actual use case in practice.
- *Lateral halves*: Images of two lateral halves of a single 18ss tailbud image. Since the point clouds do not overlap, any correspondence between the points in Lateral halves are spurious.
- *AP*: Images of the anterior and posterior ends of a single 18ss tailbud image. Like Lateral halves, any correspondence found between points in AP are spurious.

In the absence of ground truth data or an alternative image registration method, to achieve a better grasp of ZebReg’s performance, we benchmarked the registration results of these three datasets onto a noise calibration curve, which we obtained from registering noise-shifted versions of the source cloud onto its original copy.

Zero-mean gaussian distributions with standard deviations ranging from 0 to 30 were sampled to construct an array of noise matrices. These noise matrices were added to the positions of the source clouds to generate an array of noise-shifted point clouds. Conceptually, each noise-shifted point cloud is an *in silico* analogue of a tailbud that differs from its idealised, identical twin in nuclei position by a prespecified level of noise. To construct the noise-calibration curve, all noise-shifted point clouds were registered against the original source cloud, which returned the values for the fitness, inlier RMSE and inlier MAE metrics. For the heavily noise-shifted point clouds, many points are classified as outliers and therefore, the inlier metrics overestimate the registration quality by omitting these points. To correct this, we scaled the inlier RMSE and inlier MAE metrics by the corresponding fitness and plotted the scaled inlier RMSE and scaled inlier MAE values instead. Comparing our results (Figure S9E-F), we conclude that ZebReg’s registration of Test Sample outperforms the Lateral halves and AP datasets and returns acceptable fitness, inlier RMSE and MAE scores in its alignment of the Test Sample dataset.

##### Validation of ZebReg against HCR data

To assess whether the imputation procedure alters the gene expression distributions, we constructed Q-Q plots of the original and imputed gene expression distributions of 12 genes at 18ss (Figure S10A). For the purposes of the comparison of Q-Q plots, we also analysed the expression of four additional genes (wnt8a, thbs2, id3 and depdc7a) that were not used in constructing the composite maps.

We also assessed the extent to which ZebReg maintains the quantitative relationships between genes in the NMps by comparing the pairwise linear correlations of imputed genes with the original correlations from the HCR datasets (Figure S10B). A total of 17 gene pairs were compared. As a measure of how close the original and imputed correlations are to each other, we computed the minimum difference between the imputed correlation and the associated original correlations (Figure S10C). The minimum difference was computed by calculating the differences between the correlations obtained from the HCR images and the correlation from the composite map, and then taking the minimum value of the differences.

##### Construction of in silico composite maps

To construct the composite maps for each stage, we first selected images across different samples to be used for the imputation. Each image consists of *sox2* and *tbxta* stained alongside one or two additional genes and belongs to an image group. Specifically, there are a total of five image groups that correspond to particular HCR experiments (Table S2): STT (*sox2, tbxta, tcf*), STHC (*sox2, tbxta, hes6, cdh6*), STSC (*sox2, tbxta, sp5l, cdh6*), STTC (*sox2, tbxta, tagln3b, cdh6*) and STZC (*sox2, tbxta, znf703, cdh6*). For each of the three composite maps (18ss, 24ss, 28ss), five images of the same stage, one from each of the five image groups, were mapped onto a chosen target image using sox2 as the common colour channel for cICP alignment (Figure S8). These six images for each composite map were chosen to best reflect the number and spatial distributions of the *in silico* NMps in the resultant composite maps. In summary, each composite map combines information across six images (one for the target image and five for the source images) to generate an eight-dimensional (*sox2, tbxta, cdh6, hes6, sp5l, tagln3b, tcf, znf703*) point cloud image.

#### Analysis of the composite maps

##### Identification of in silico NMps

Following the construction of the composite map with ZebReg, imputed intensity values of all genes below the 0.7 quantile threshold were set to 0 and rescaled by min-max normalization (Figure S12A). Amongst the *sox2*+*tbxta*+ cells that were identified in the composite map, most were found within the approximate NMp spatial regions (Figure S12Bi-ii). Of the *sox2*+*tbxta*+ points that reside outside of the NMp regions and are thus excluded from being NMps, they fall into two groups (Figure S12Bii). The first group corresponds to the hypochord cells that constitute the bulk of these *sox2*+*tbxta*+ non-NMps in the 18ss (47/146: 32%), 24ss (43/203: 21%) and 28ss (29/56: 52%) composite maps. The second group of cells are found in the 18ss composite map only (17/146 : 11%) and are a small population of aberrant cells that have likely arisen from technical errors in either ZebReg’s alignment or mapping procedure. These cells flank the hypochord and floor plate and thus may have been mistakenly assigned above-background levels of *sox2* due to their proximity to these two *sox2*-expressing structures. After the removal of these two groups of cells, the resultant 78, 183 and 27 *in silico sox2*+*tbxta*+ cells in the 18ss, 24ss and 28ss composite maps are defined as the *in silico* composite map NMps. (Figure S12Biii)

##### Construction of the Neural-Mesodermal (NM) Index

The neural-mesodermal index, *NM*_*j*_ for the *j*^*th*^ cell is defined as:

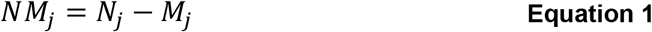

where *NM*_*j*_, *N*_*j*_, *M*_*j*_ are the neural-mesodermal index, neural index and mesodermal index of the *j*^*th*^ cell, respectively, and *j* = 1,2, …, *C* for a total of *C* NMps.

The neural index, *N*_*j*_, for the *j*^*th*^ cell is defined as:

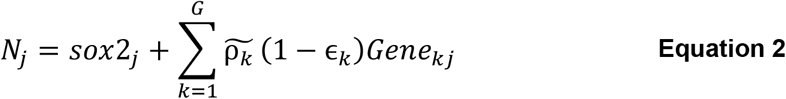

where *Gene*_*kj*_ is the min-max normalised expression intensity of the *k*^*th*^ gene in the *j*^*th*^ cell; 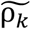 is the median of the Pearson’s correlation coefficients of *Gene*_*k*_ and *sox2*, computed from the NMps segmented from all the HCR images of the same somite stage; *ϵ*_*k*_ is the interquartile range of *Gene*_*k*_’s correlation coefficients. The (1 − ***ϵ***_*k*_) term penalises *Gene*_*k*_’s contribution to the neural index if it displays large variability in its correlation coefficients between all the HCR images of that somite stage. The summation is applied to all the genes, *G*, minus *tbxta* and *sox2*. The total number of genes is *G* + 2.

The mesodermal index, *M*, is defined symmetrically but with *tbxta* replacing *sox2* and the correlation coefficients calculated with respect to *tbxta* instead. We further verified that the NM index provides a sensible summary of the NMp’s neural/mesodermal potential (Figure S13B-D) and is not systematically biased towards either neural or mesodermal indices (Figure S13E-E’).

##### Construction of the NMp probability map

Tailbud images (source images) were aligned to an arbitrarily chosen target image tailbud. The NMp nuclei in the source images were pre-segmented prior to alignment in Imaris (Bitplane) and hence, it is possible to keep track of the number of times each target cell receives a mapping from a source NMp cell. Target cells with a large count number is assigned a high probability of being an NMp. For visualisation purposes, in my probability maps, we displayed only target cells with a minimum count number of 2 for each probability map (Figure S14).

##### Estimation of the standard error of empirical entropy

The standard error was estimated by the leave-one jackknife resampling method and is implemented using the R *bootstrap* package (Efron and Tibshirani, 1993; Wiesner et al., 2017). In this method, the entropy was repeatedly estimated but with one of the data points randomly removed during each computation.

#### Dynamical systems modelling of a toggle switch model

##### Deterministic model formulation

The deterministic model consists of two continuous ODEs describing the rate of change of protein concentrations (Figure 4E, Figure S14). As shown in Equation 3 and Equation 4, the regulatory interactions between genes are summarized by Hill functions, whereas the decay rate of each product is set to be linearly proportional to the protein concentration.

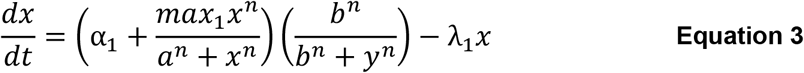

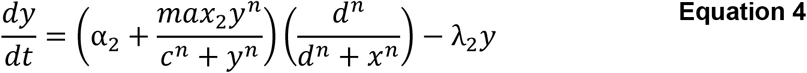

where:

- *x* and *y* are the concentrations of the two proteins X and Y.
- *α*_1_ and *α*_2_ are the rates of production of gene X/Y, in the absence of activation by X/Y.
- *n* is the Hill coefficient and is set to 4 for all simulations.
- *a* and *c* are the concentrations for the half-maximal activation by genes X/Y on genes X/Y. The inverses, 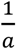 and 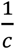 represents the efficiency of the activator in other equivalent formulations (Goutsias & Kim, 2004).
- *b* and *d* are the concentrations for the half-maximal repression by genes X/Y on genes Y/X. The inverses, 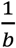 and 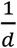 represents the efficiency of the repressor in other equivalent formulations.
- λ_1_ and λ_2_ are the protein degradation rates.
- *max*_1_ and *max*_2_ are the maximum rates of protein production by activators X / Y. To see this, take 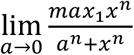.

##### Classification of stability of steady states

The stability of the steady states was determined by linear stability analysis (Strogatz, 2000). Briefly, the Jacobian matrix for the ordinary differential equations (Equation 3 and Equation 4) was constructed to obtain an approximate description of the phase portrait around the fixed points. By neglecting the quadratic and higher order terms in the Taylor series expansion, one can describe the dynamics of the linearised system with the Jacobian matrix. To assess the stability of the fixed points, we computed the determinant of the Jacobian at each equilibrium point. A negative determinant corresponds to a saddle, while a positive determinant corresponds to a sink (stable node), source (unstable node) or center. In our model, all equilibria with positive determinants correspond to sinks.

##### Stochastic toggle switch model formulation

As per the deterministic formulations, the rate of production of X and Y can be written as:

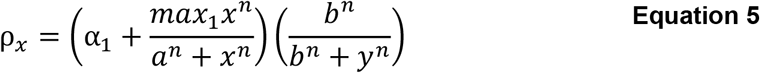

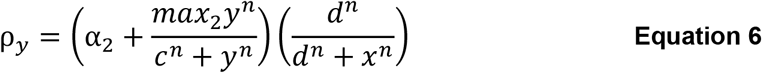

The chemical langevin equation (CLE) approximation for the process leads to the following white noise form of the Langevin equation:

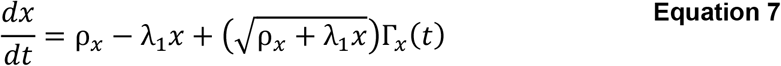

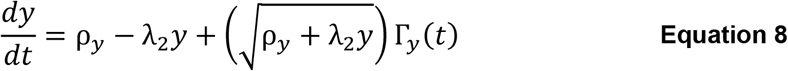

Here, the Γ_*i*_(*t*) terms correspond to Gaussian white noises, which are formally defined as (Gillespie, 1996, 2000):

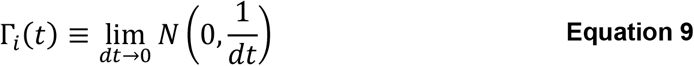

The two averaged properties of Gaussian white noise processes are:

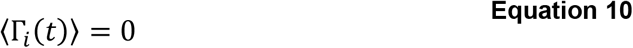

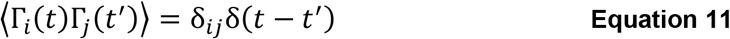

The first property indicates that the white noise process has zero mean. In the second, the first delta function is Kronecker’s delta and the second is Dirac’s delta, which indicates that white noise processes are statistically independent and that the white noise process is temporally uncorrelated.

##### Numerical simulation with the Euler-Maruyama method

The standard form of the Langevin equation is written as the following difference equation, where Ω is the volume parameter (Gillespie, 2000):

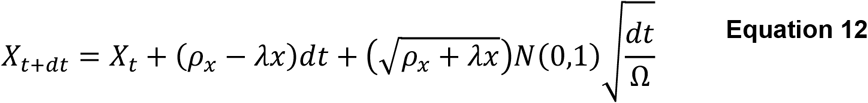

In the limit Ω → ∞, we obtain the deterministic result.

Numerical simulation of the CLE was carried out with the Euler-Maruyama algorithm, which has the following numerical scheme that approximates the infinitesimal *dt* with finite Δ*t* (Wilkinson, 2020):

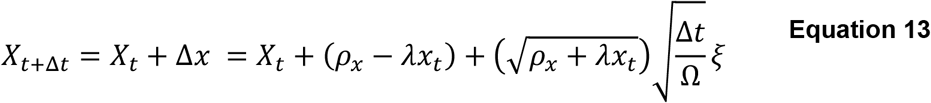

If the molecule count number dips below zero, it is automatically set back to zero to prevent the molecule numbers from assuming negative values.

##### Modelling non-autonomy via a stepwise approximation approach

The incorporation of non-autonomy was carried out following the work of Verd and colleagues (Verd et al., 2014), where the parameter value is kept constant over a short time step and incremented at regular intervals.

#### Quantification and Statistical Analysis

For all boxplots in

Figure **2**, the lower and upper hinges correspond to the first and third quartiles. In addition, the upper whisker extends from the hinge to the largest value no further than 1.5 times the interquartile range. Outlier samples are coloured in red. Wilcoxon-Mann-Whitney unpaired two-sample test ****: p-value < 0.0001; * p-value < 0.01; ns = not significant.

