## Supplementary Figure for "Zebrafish Neuromesodermal Progenitors Undergo a Critical State Transition *in vivo*"

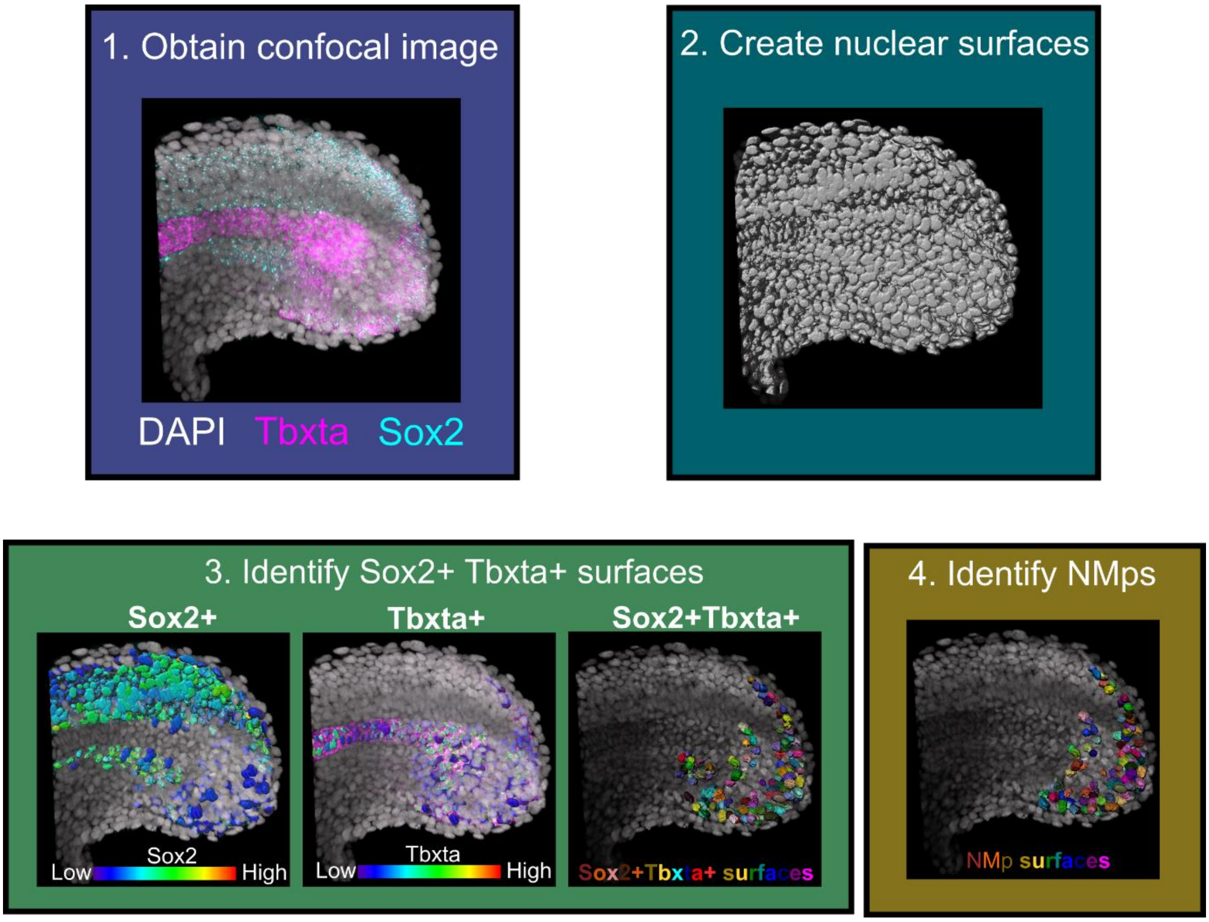


**Figure S1: Segmentation of NMps from confocal images with Imaris, related to STAR methods.**

After acquiring the confocal image (1), general surfaces were created (2) on the DAPI channel of the entire image with the following key parameters: Surfaces Detail = 0.300, Split touching Objects (Region Growing) = Enabled, Seed Points Diameter = 3. When classifying the seed points for these general surfaces, all points were selected by default unless outlier spots were present. If so, these outlier spots were removed using the Quality filter.

Next, *tbxta*+ *sox2*+ surfaces were selected by filtering from the general surfaces (3). Specifically, surfaces with *tbxta* mean intensity and *sox2* max intensity above both of their respective automatic lower thresholds were selected. These surfaces represent nuclei that co-express above background levels of *sox2* and *tbxta*. The max intensity threshold was chosen for *sox2* as there is an increased background autofluorescence in the 488 nm imaging channel, and the max intensity threshold was found to be more accurate in selecting the *sox2*+ surfaces than the mean intensity threshold. We then verified the suitability of the intensity thresholds by comparing to their known expression domains.

Finally, *tbxta*+*sox2*+ surfaces were created and surfaces in the hypochord region were excluded (4). The remaining *tbxta*+*sox2*+ surfaces were classified as NMp surfaces. The mean intensities of *sox2* and *tbxta* were exported from these NMp surfaces.

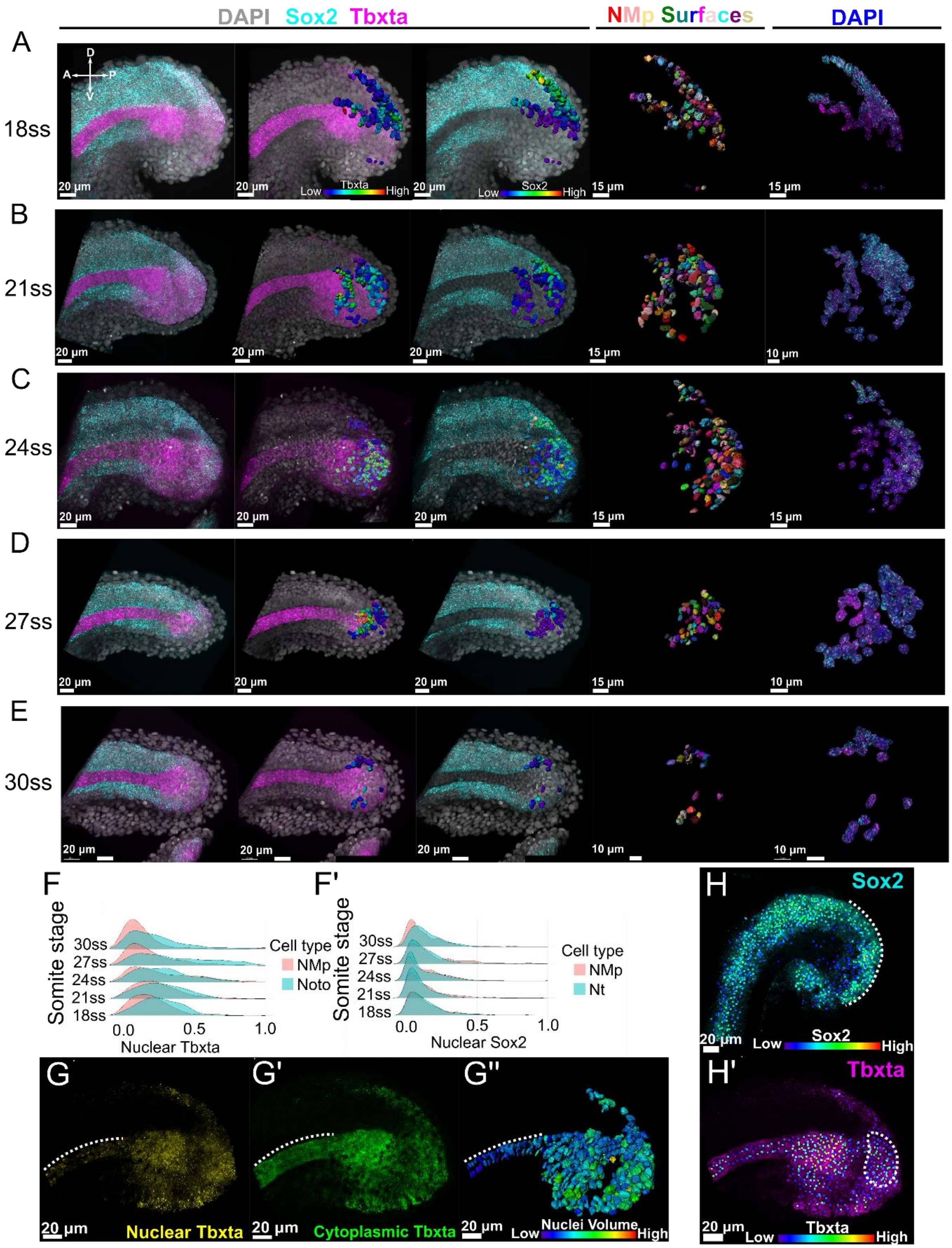


#### Figure S2: *Sox2* and *Tbxta* expression in the NMps, neural tube and notochord populations, related to Figure 1.

(A - E) Time-course HCR experiment of the zebrafish tailbud NMps from 18ss to 30ss. Each panel depicts the imaging channels and segmented NMp nuclei for a single sample. (From left to right) 3D composite image of *sox2*, *tbxta* and DAPI. (Second column) NMp surfaces coloured by *tbxta* intensity, shown alongside *tbxta* expression. (Third column) NMp surfaces coloured by *sox2* intensity, shown alongside *sox2* expression. (Fourth column) NMps depicted as rainbow surfaces. (Fifth column) Magnified images of the NMps masking the expression of *sox2* and *tbxta* outside of the surfaces.

(F-F') Ridge plots comparing the (F) nuclear *tbxta* distributions of the NMp and the notochord populations. (F') nuclear *sox2* distributions of the NMp and caudal neural tube populations

(G-G'') (G) Nuclear expression of *tbxta*. (G') Cytoplasmic expression of *tbxta*. (G'') *tbxta*-positive nuclear surfaces coloured by nuclear volume. Dashed lines refer to the notochord region which appears to have higher levels of cytoplasmic *tbxta* relative to nuclear *tbxta*.

(H-H') (H) *Sox2* positive nuclei as points coloured by *sox2* intensity (H') *Tbxta* positive nuclei as points coloured by *tbxta* intensity.

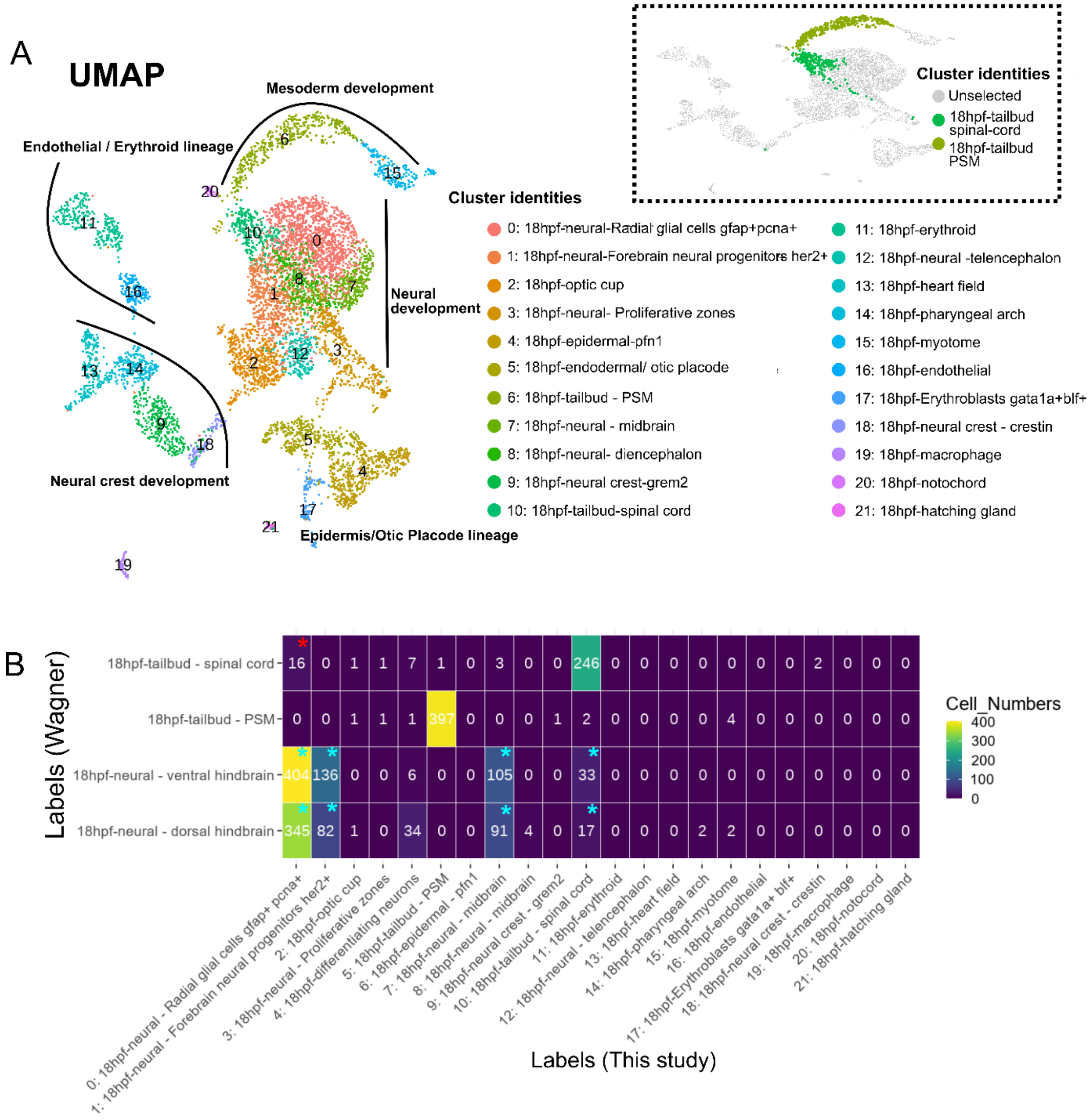


**Figure S3: UMAP visualisation of the 18hpf dataset and distribution of Wagner and colleagues' labels across clusters, related to Figure 2.**

(A) UMAP visualisation of the 6,959 zebrafish cells at 18hpf colour coded according to assigned cluster. The inset at the top-right highlights the tailbud spinal cord and PSM clusters.

(B) Heatmap displaying the distribution of 4 of Wagner and colleagues' labels across our Louvain tailbud clusters. The Wagner tailbud labels are highly concordant with our tailbud labels; with the only notable exception being that 16 of the radial glial cells (Louvain cluster 0) fall into Wagner's 18hpf-tailbud - spinal cord cluster (red asterisk). The similarity in the tailbud cluster annotations contrasts with the neural hindbrain clusters (cyan asterisks) which consist of cells from our Louvain clusters 0, 1, 7 and 10.

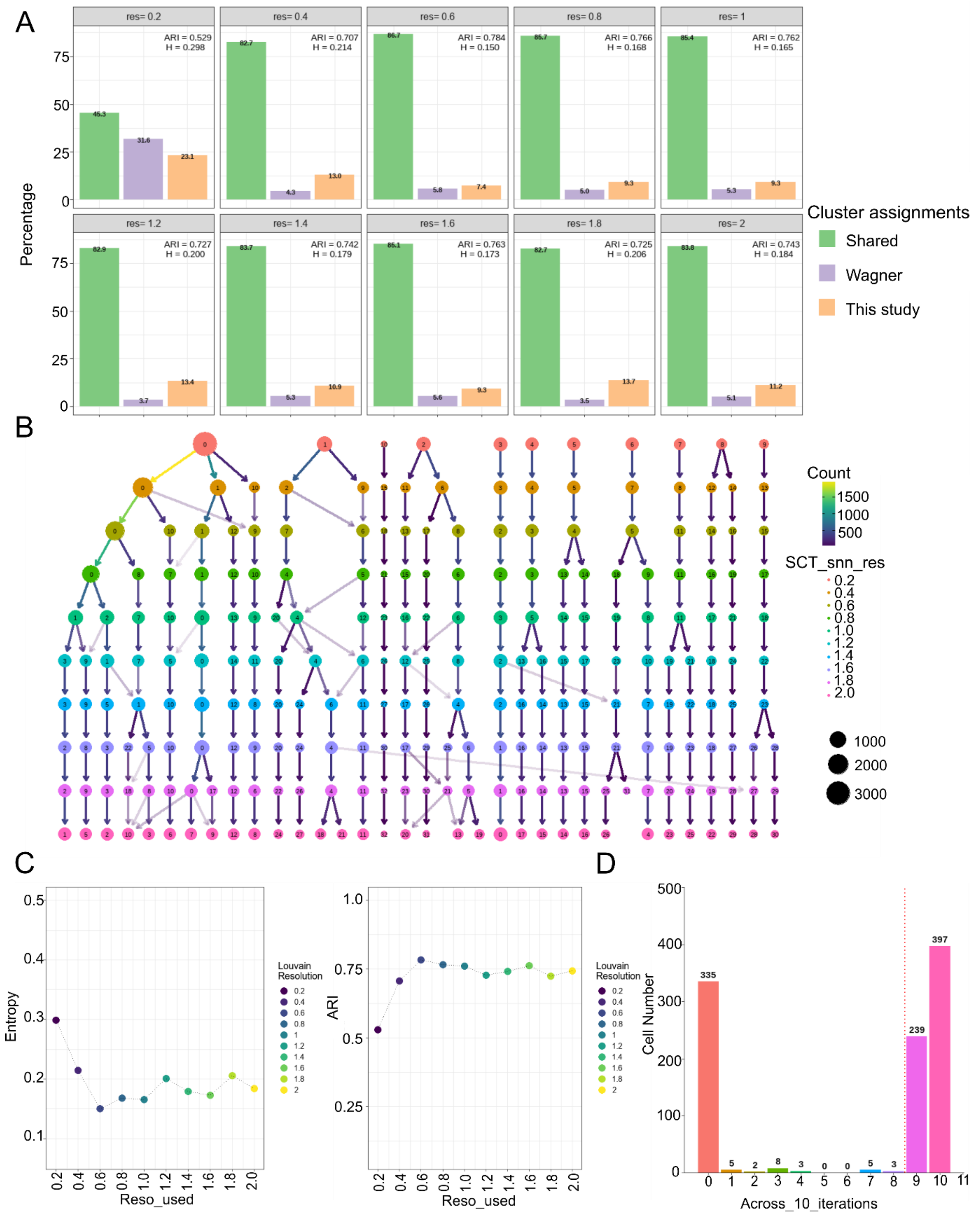


**Figure S4: Optimizing parameters for Louvain clustering and tailbud cell selection, related to STAR methods.**

(A) Faceted bar plots assessing the distribution of cell labels across ten (0.2 - 2.0) different Louvain clustering resolutions. Cells in the shared category were identified by Wagner and colleagues to be in the tailbud and are also assigned to the tailbud spinal cord or PSM clusters in this study. Cells in the

‘Wagner’ category were identified by the Wagner and colleagues to be in the tailbud but are not assigned to the Louvain tailbud clusters in this study. Cells in the ‘This study’ category are assigned to the Louvain tailbud clusters but were not identified as belonging to the tailbud by Wagner and colleagues. ARI: Adjusted rand index; H: Entropy.

(B) Clustering tree displaying how cells move across clusters as the Louvain resolution increases (greater number of clusters).

(C) Line plot displaying the change in clustering entropy (Left) and ARI for the tailbud clusters at different resolutions, using Wagner and colleagues’ labels as ground truth.

(D) Bar plot displaying the number of cells that are assigned to the shared category across all 10 iterations of the Louvain clustering algorithm, with a different resolution parameter for each iteration. For instance, 239 cells were assigned to the shared category in 9 out of the 10 iterations. The dotted red line indicates the threshold above which the cells are identified as tailbud cells and retained for downstream analysis.

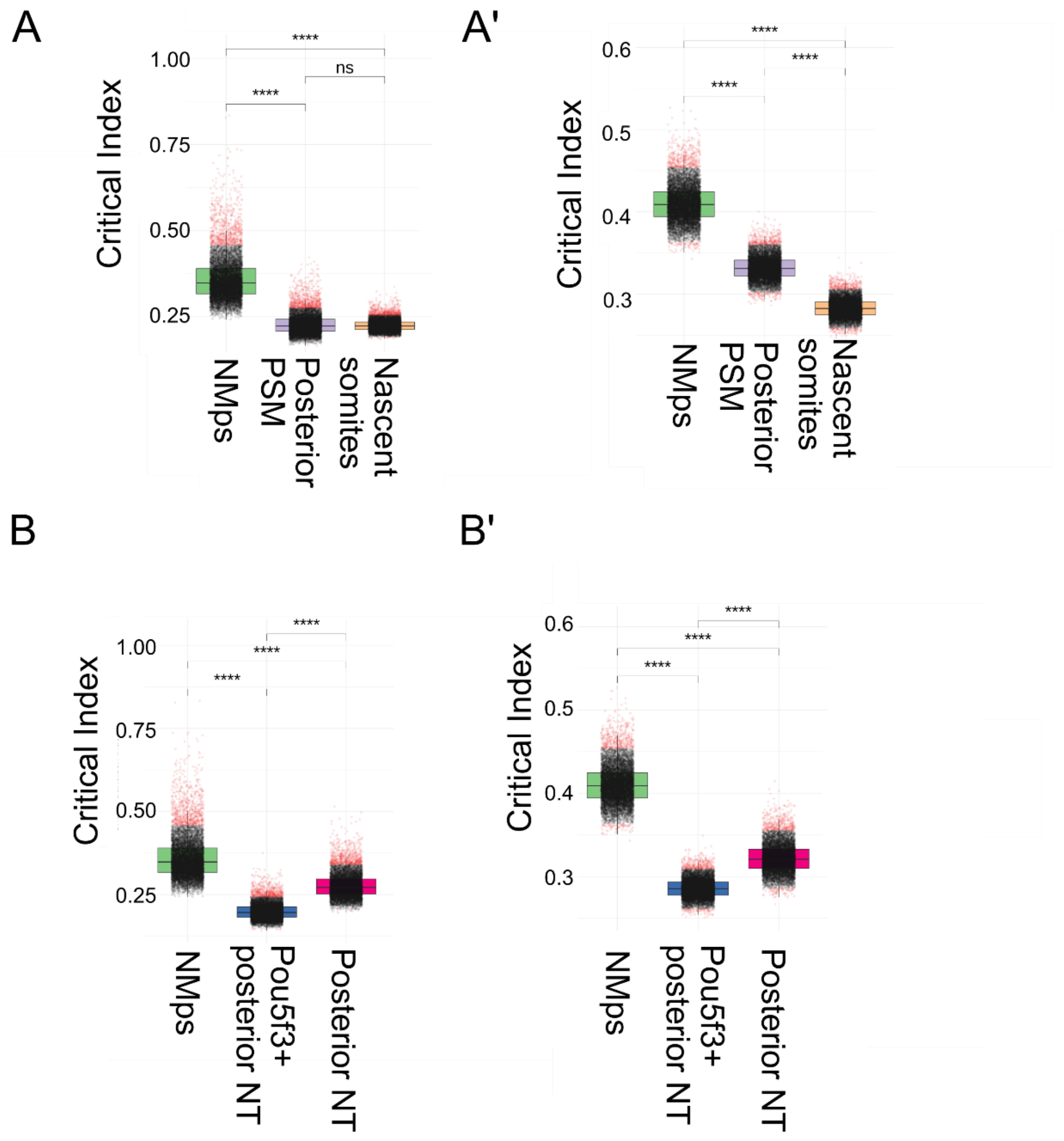


**Figure S5: Statistical robustness of the critical index, related to Figure 2.**

(A - A') Critical index calculation along the mesodermal branch. (A) Instead of using all the marker genes to compute the index for each cluster, 100 marker genes were randomly chosen each time, and the procedure was repeated for 10,000 times. (A') Instead of randomly sampling 200 cells from each cluster with replacement, 45 cells were randomly sampled from each cluster with replacement each time, and the procedure was repeated for 10,000 times.

(B - B') Similar calculations as (A - A') but for the neural branch.

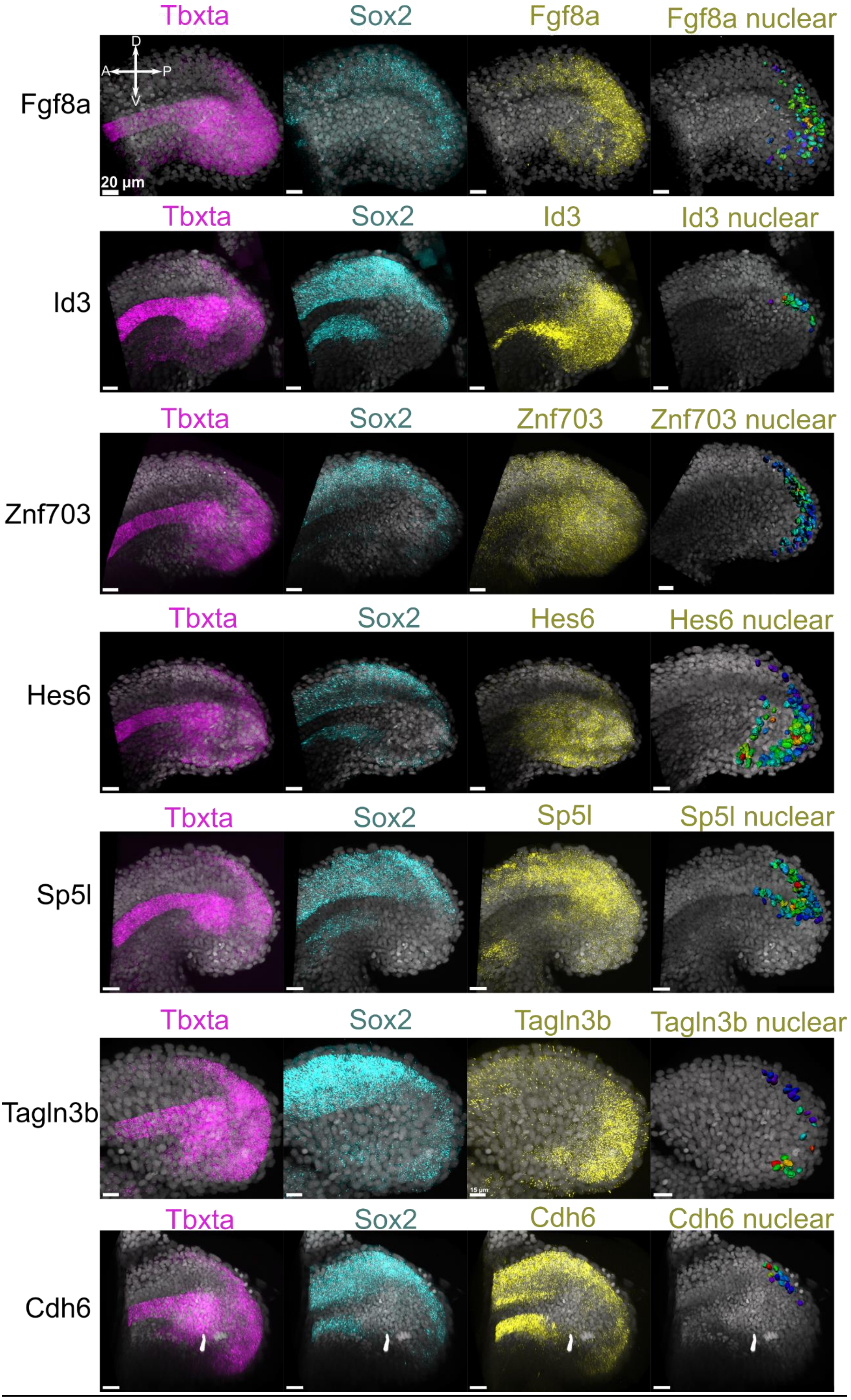
Tbxta
Sox2
Fgf8a
Fgf8a nuclear
Fgf8a
D
A
P
V
20 µm
Tbxta
Sox2
Id3
Id3 nuclear
Id3
Tbxta
Sox2
Znf703
Znf703 nuclear
Znf703
Tbxta
Sox2
Hes6
Hes6 nuclear
Hes6
Tbxta
Sox2
Sp5l
Sp5l nuclear
Sp5l
Tbxta
Sox2
Tagln3b
Tagln3b nuclear
Tagln3b
Tbxta
Sox2
Cdh6
Cdh6 nuclear
Cdh6

**Figure S6: Validation of 7 marker genes identified from the scRNA-seq analyses, related to Figure 2.**

Expression of *tbxta*, *sox2* and a selected NMp marker gene in the zebrafish tailbud (For full list of marker genes, see Table S3). (From left to right) *Tbxta* expression. (Second column) *Sox2* expression. (Third column) Selected marker gene expression. (Fourth column) Triply positive surfaces (*sox2*, *tbxta* and marker gene) coloured by the gene expression levels of the selected marker gene. All scalebars are 20 μm. DAPI is in grey.

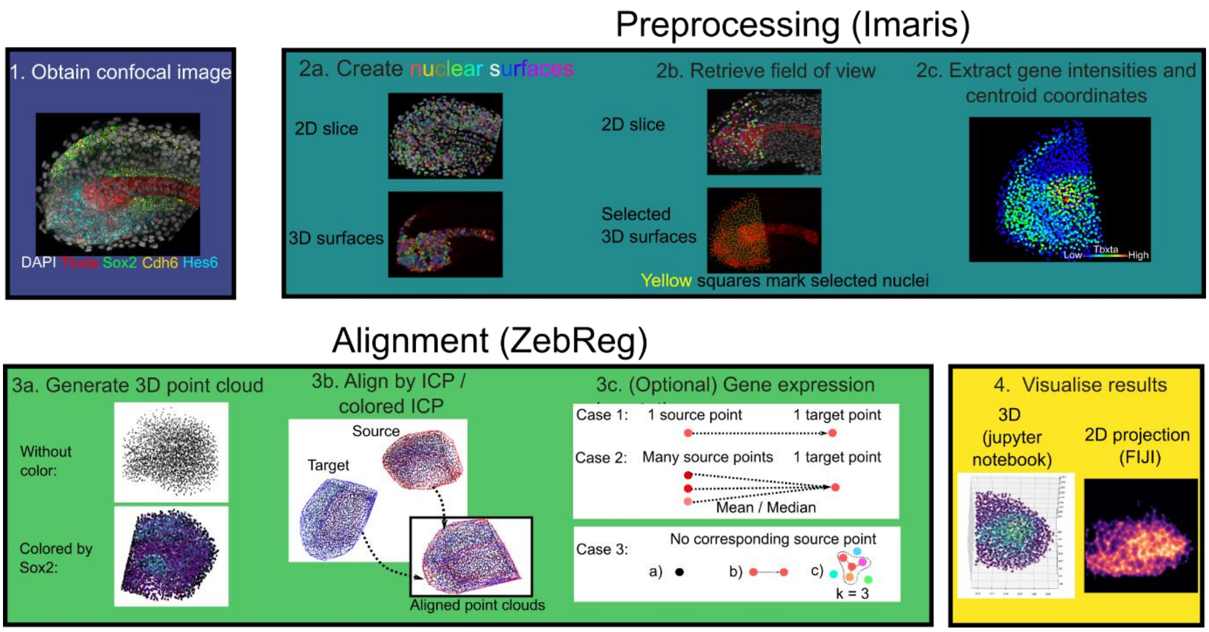


**Figure S7: Four stages of the ZebReg image registration pipeline, related to STAR Methods.**

In the data acquisition stage, zebrafish tailbuds were stained with HCR and counterstained with DAPI, which label the mRNA and nuclei respectively. Tailbuds were imaged by confocal fluorescence microscopy under a 40x objective. Next, the image is preprocessed in Imaris (bitplane). To align the tailbud images in ZebReg, ZebReg uses the 3D centroid coordinates as the position coordinates for the point cloud and can also store the expression intensity values in various colour channels in each point. Images can be aligned by either the Iterative Closest Point (ICP) or coloured ICP (cICP) algorithms. Gene expression intensities can also be imputed onto the target point cloud if present in the source cloud (For more detail, see *imputation of gene expression intensities* in Methods). Following the mapping, one can visualise the coloured points within Jupyter Notebook or export the image stack.

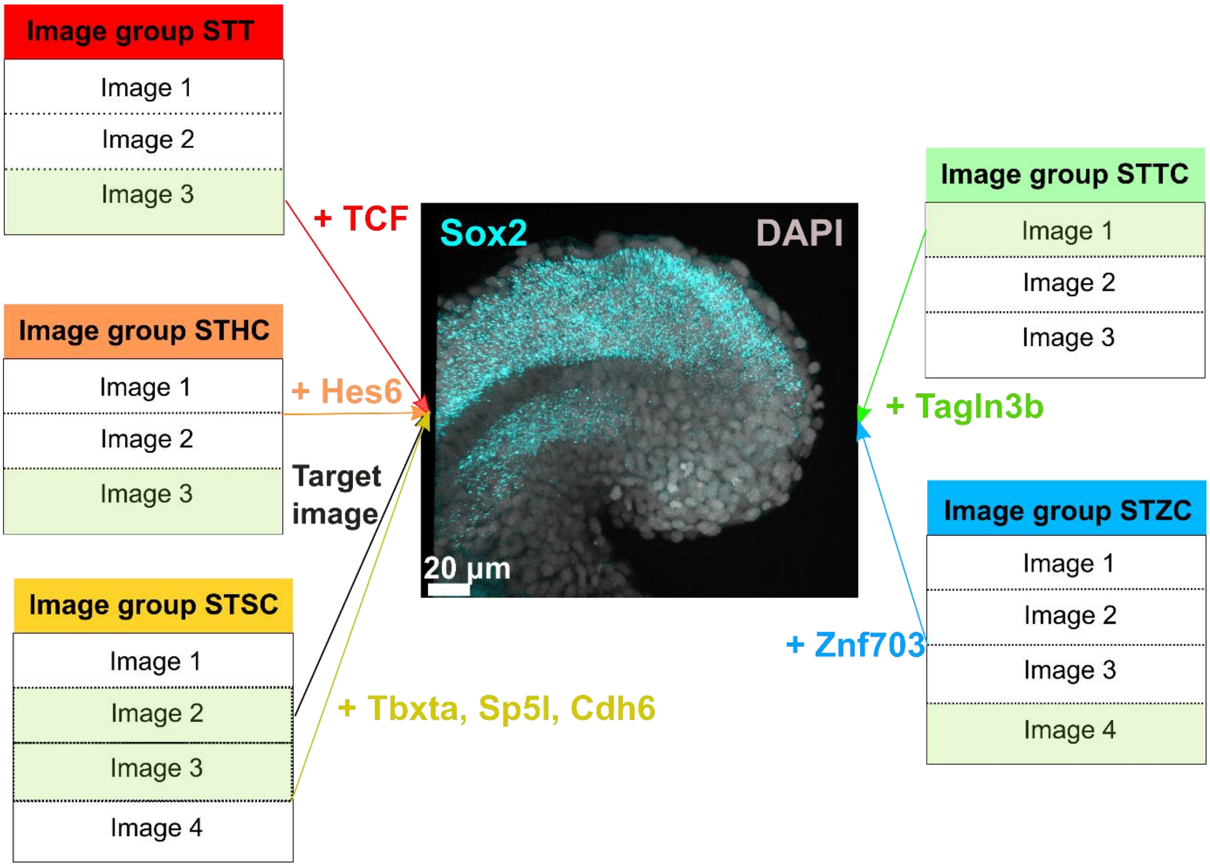


**Figure S8: Construction of the composite map, related to STAR Methods.**

The target image is shown in the middle of the panel and corresponds to Image 2 from the STSC image group, with its *sox2* expression in cyan. The gene(s) selected for imputation onto the target image from each image group are indicated.

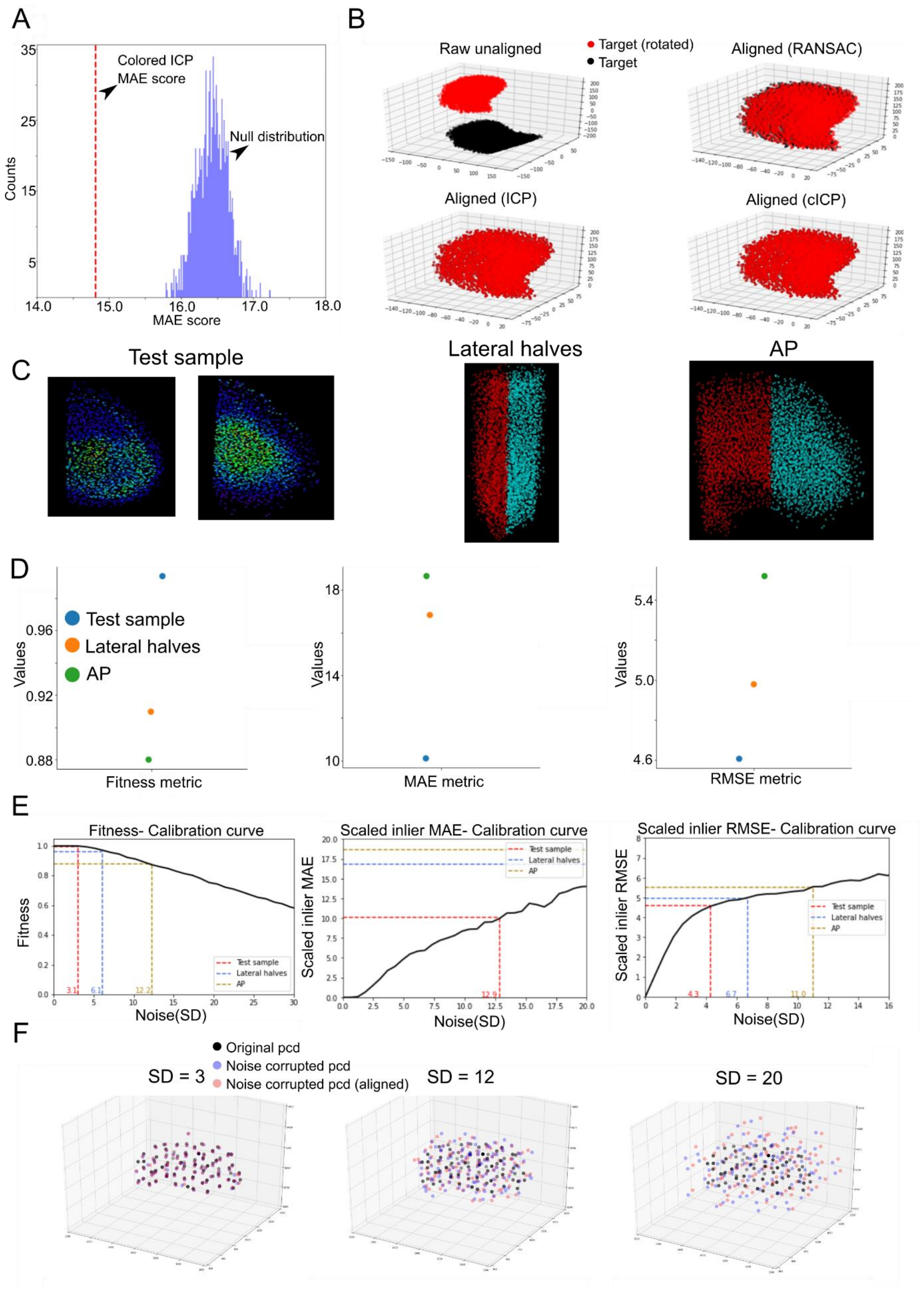
A
Colored ICP MAE score
Null distribution
Counts
MAE score
B
Raw unaligned
Target (rotated)
Target
Aligned (RANSAC)
Aligned (ICP)
Aligned (cICP)
C
Test sample
Lateral halves
AP
D
Test sample
Lateral halves
AP
Values
Fitness metric
MAE metric
RMSE metric
E
Fitness- Calibration curve
Scaled inlier MAE- Calibration curve
Scaled inlier RMSE- Calibration curve
Fitness
Scaled inlier MAE
Scaled inlier RMSE
Noise(SD)
F
Original pcd
Noise corrupted pcd
Noise corrupted pcd (aligned)
SD = 3
SD = 12
SD = 20

**Figure S9: In silico validation of ZebReg, related to STAR Methods.**

(A) Comparison of the cICP MAE score against the corresponding null distribution. The empirical cICP MAE score of 14.81 (dotted red lines), which was obtained from registering the source and target point clouds, lies outside of the 95% CI (16.00, 16.85) of the null distribution. Therefore, there is sufficient evidence to support the claim that cICP reduces the colour intensity residuals between point clouds.

(B) Alignment of a source and target point cloud by different algorithms. In this trivial example, the target cloud is aligned against a linearly transformed version of itself. Both ICP and cICP gave a perfect alignment of the point clouds, whereas RANSAC's performance was inferior and provided only a rough alignment. RANSAC: Random Sample Consensus ICP: Iterative Closest Point; cICP: coloured ICP.

(C) Images of the Test sample, Lateral halves and AP datasets.

(D) Performance of the Test sample, Lateral halves and AP datasets evaluated on three metrics. For the fitness metric, the higher the value, the better the performance. For the MAE and inlier RMSE metrics, the lower the value, the better the performance. The Test sample registration outperforms the Lateral halves and AP registration.

(E) Calibration curves for the fitness, MAE and inlier RMSE metrics. The bolded black lines are the calibration curves which correspond to the values of the metrics when the registered image is subjected to varying levels of noise-corruption (See *In silico validation of ZebReg* section in *STAR Methods)*. With increasing noise standard deviation (SD), the performance of cICP degrades along all metrics. From the perspective of the MAE metric (Middle), registering the Test sample dataset is akin to registering an image onto its associated noise corrupted copy with a noise SD of 12.9.

(F) 3D visualisation of the original point cloud, alongside the unaligned and aligned noise-corrupted point clouds at increasing standard deviation of noise. At noise SD = 3 and SD = 12, the point clouds still appear relatively well-aligned to each other but at noise SD = 20, they are poorly aligned.

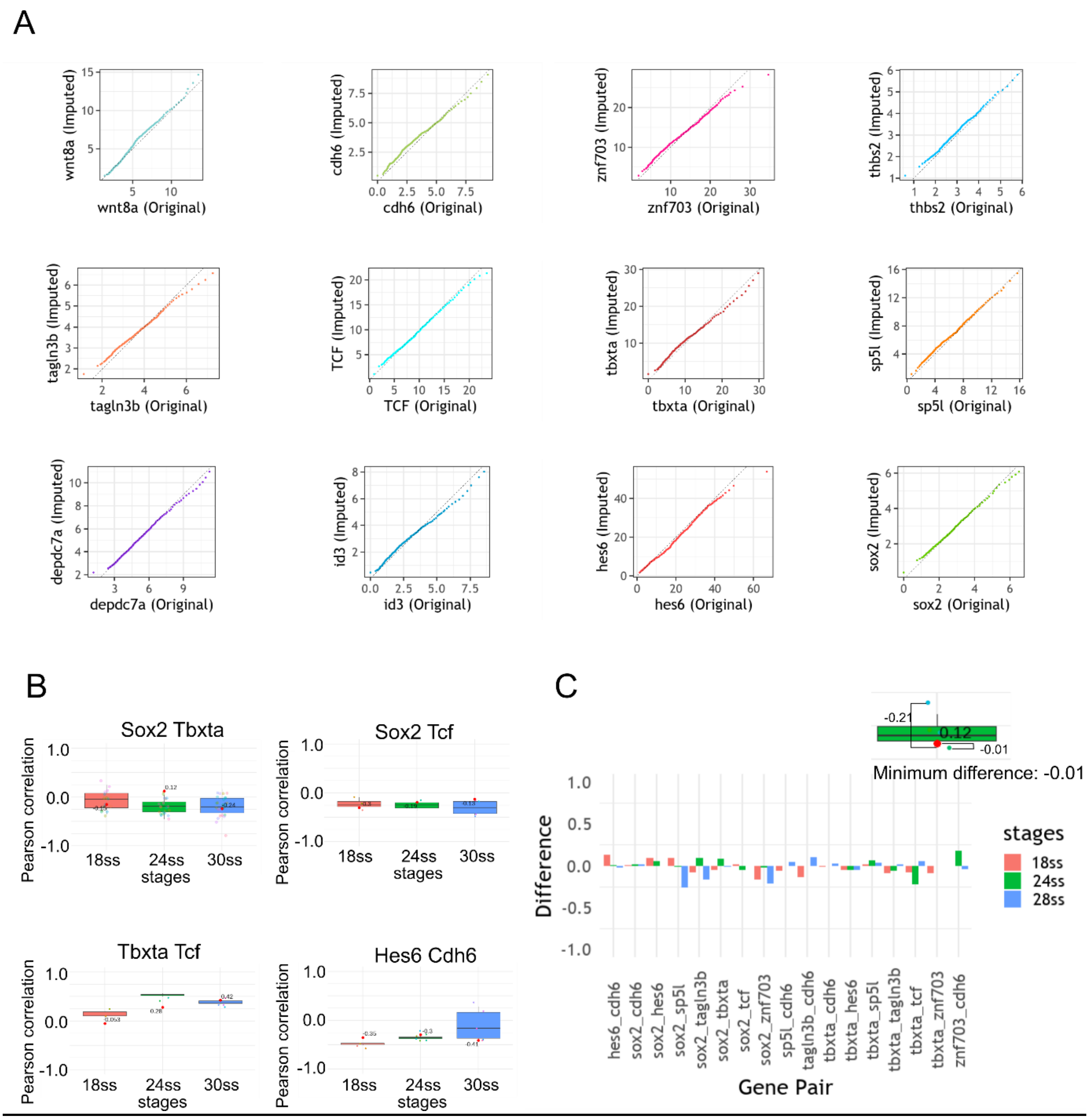


**Figure S10: Validation of ZebReg against HCR data, related to STAR Methods.**

(A) Q-Q plots of the original and imputed gene expression distributions. Most of the points lie approximately along the y=x line. Points with significant deviation from the line correspond to values with extreme percentile scores. The averaging procedure reduces the variance of the original distribution, affecting the expression values at the tails more significantly, but it does not have a significant impact on the distribution otherwise.

(B) Boxplots showing the distributions of the pairwise correlations between genes obtained from the segmentation of NMps in HCR images. Red points indicate the correlations obtained from the *in silico* composite maps.

(C) Minimum difference plot of all 17 gene pairs. The inset depicts the computation of the minimum difference. Across the 17 gene pairs in all three stages, the minimum differences between the original and imputed correlation values lie within the range of -0.27 to 0.18. Additionally, these differences do not show a positive or negative bias, nor is there any apparent relationship between the stages and the magnitude of the differences. The absence of systematic errors suggests that the registration quality is reasonably consistent across the three stages.

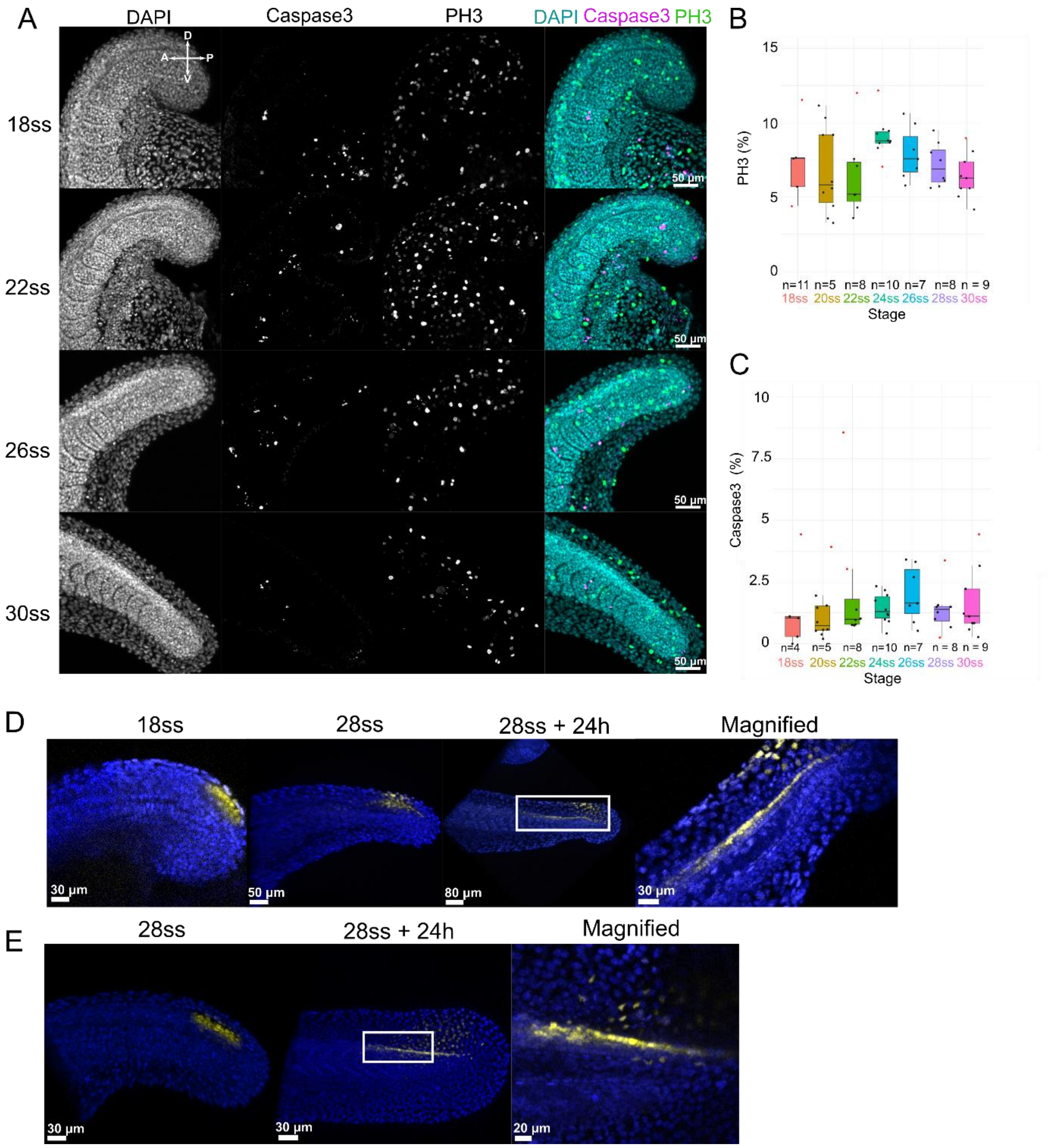


**Figure S11: Characterisation of proliferation, apoptosis, and migration dynamics in the NMp population, related to Figure 5.**

(A) Zebrafish tails from 18ss to 30ss were stained and detected for Caspase3 and PH3 by immunofluorescence to estimate the levels of apoptosis and cell division respectively. Nuclei were stained with DAPI. All tails are orientated towards the right.

(B - C) Percentage of (B) PH3-positive nuclei and (C) Caspase3-positive nuclei from 18ss until 30ss with an interval of two somites between stages. The levels of proliferation and apoptosis in the tailbud remain low from 18ss to 30ss. *n* refers to the total number of samples analysed for each stage and is indicated above the x-axis.

(D) Photolabels (yellow) made in the dorsal posterior wall (PW) at 18ss was followed until 28ss. By this point (28ss), all the photolabelled cells reside in the fin mesenchyme and the dorsal neural tube which implies that all the NMps in the dorsal PW move anteriorly into the dorsal NT. When the tailbud was allowed to develop for another 24 hours (28ss+24h), the photolabels were found to have spread more anteriorly, with the anterior labels appearing more dispersed ventrally (Magnified).

(E) A similar phenomenon to (D) is observed when the dorsal PW is photolabelled at 28ss and reimaged 24h later (28ss+24h).

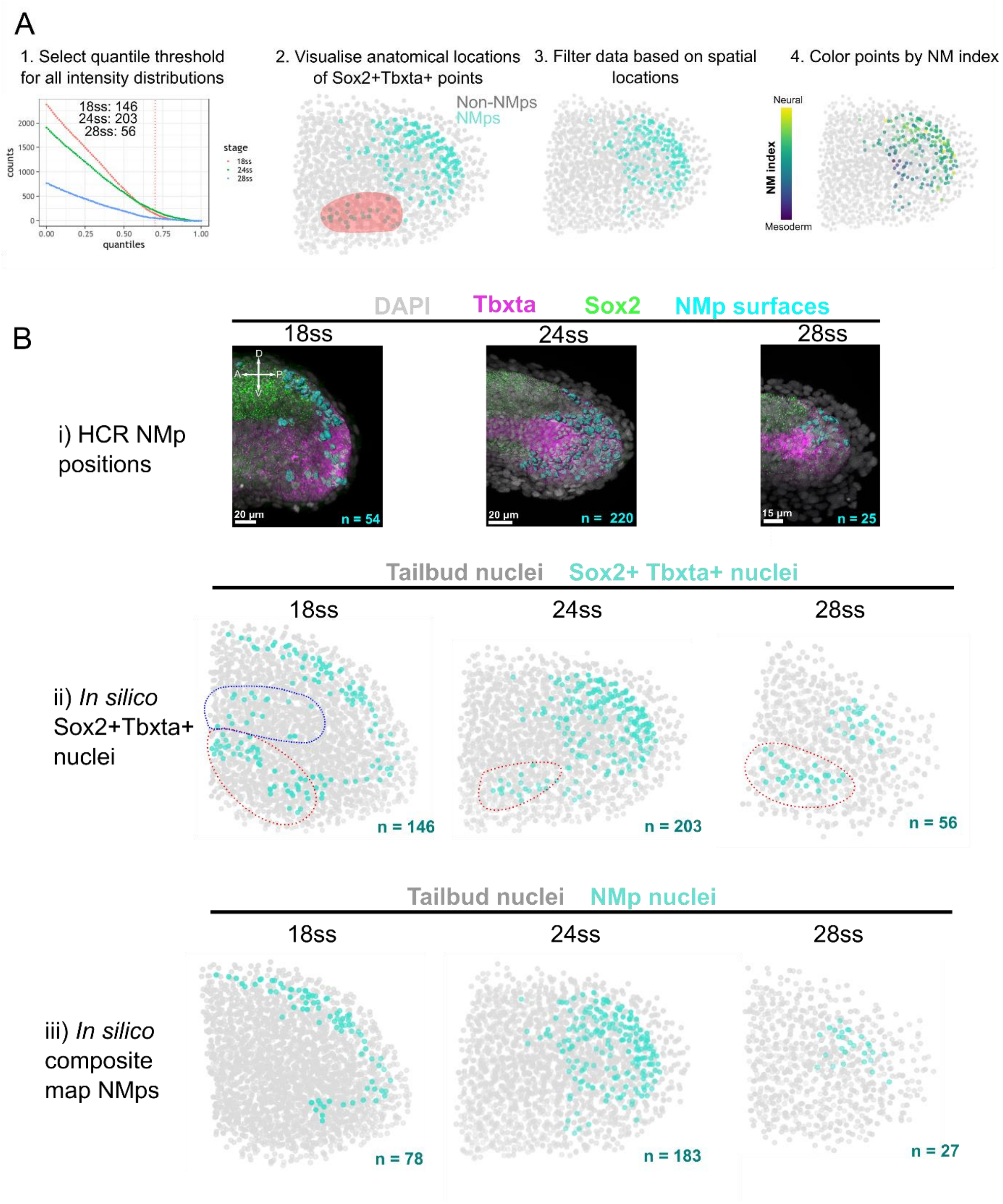


**Figure S12: ZebReg recovers the spatial positions of zebrafish NMps, related to Figure 5.**

(A) Procedure to identify the *in silico* composite map NMps.

(B) (i) HCR images of tailbuds stained for *sox2* and *tbxta* with segmented NMp nuclei represented as cyan surfaces. Note that these HCR images are not the same images that were selected for the imputation of *sox2* and *tbxta* in the composite map shown below. (ii) Point cloud representation of the tailbud from 18ss, 24ss and 28ss, with the locations of *sox2+tbxta+* nuclei coloured in cyan. Dashed blue circle highlights aberrant cells that most likely arise from registration error. Dashed red circle highlights hypochord cells. (iii) Point cloud representation of the tailbud, with the locations of in silico composite map NMps coloured in cyan. The number of identified *sox2+tbxta+*/NMp nuclei is shown at the bottom right-hand corner of the images.

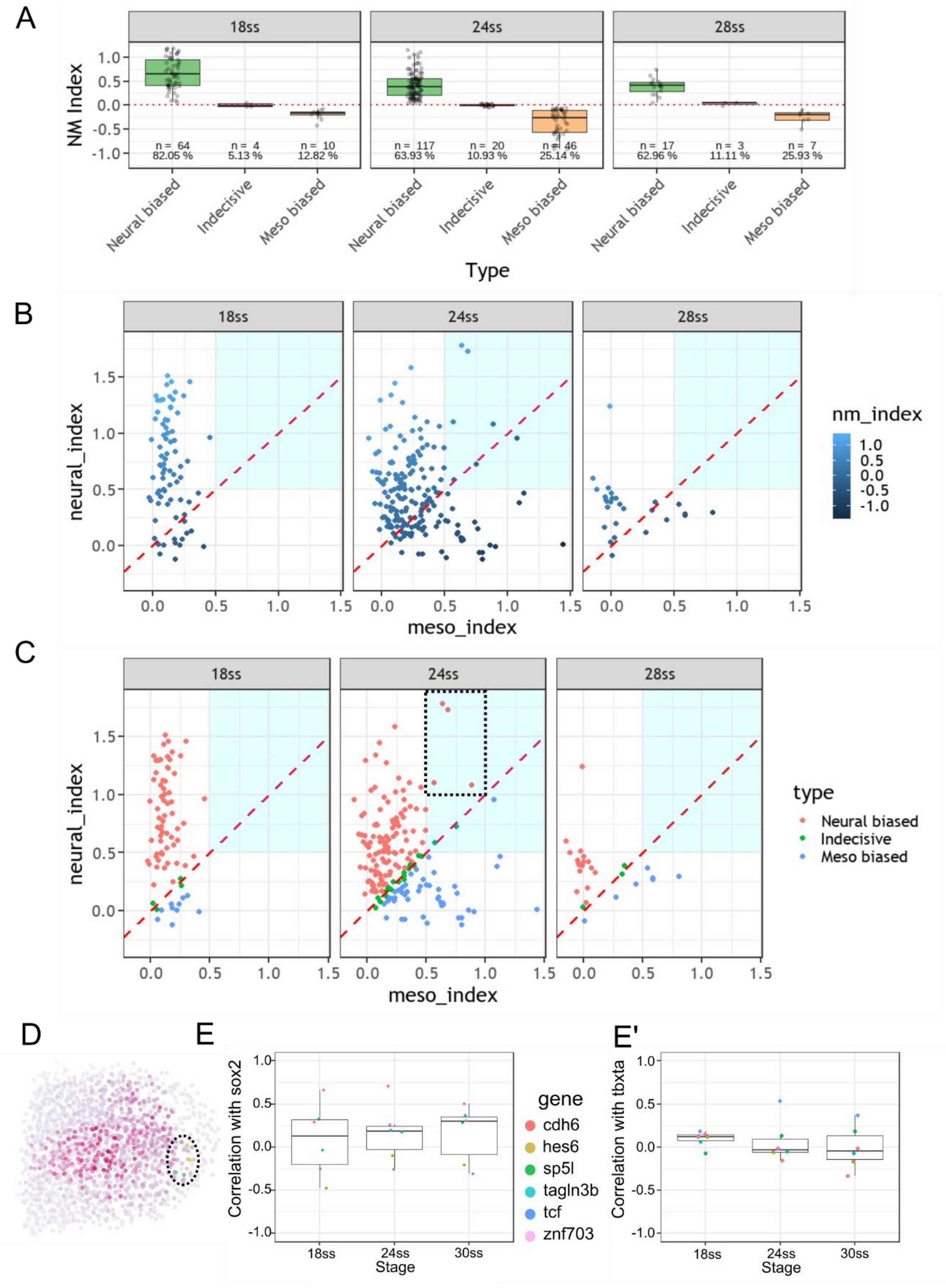


**Figure S13: Visualising the relationship between the Neural and Mesodermal indices, related to Figure 5.**

(A) Proportion of NMps in 3 different expression categories. 'Neural biased' NMps have an NM index greater than 0.05, whereas 'Meso biased' NMps have an NM index less than -0.05. NMps with an NM index between -0.05 and 0.05 are 'Indecisive'.

(B - C) In silico NMps from the composite maps are coloured according to their (B) NM index (C) type. The dashed red line represents the y=x line. Points lying on this line are Indecisive. The cyan region at the upper right quadrant highlights cells with high values of both the Neural and Mesodermal indices. The boxed region in (C) highlights neural biased cells with high values of the Mesodermal index. At 18ss, most cells have a high Neural and low Mesodermal index, confirming our inference that most cells at 18ss are neural biased. Apart from the 24ss scatterplot, there are no cells in the upper-right quadrant at the remaining stages. The upper-right quadrant demarcates a region where cells have high levels of both indices. At 24ss, 4 neural biased, 2 Indecisive and 1 mesoderm biased cell at 24ss fall in this region.

(D) Location of the four neural-biased cells in the boxed region (C) are circled. These cells, with relatively high Mesodermal indices, fall in the mesoderm-fated spatial domain and are thus also Incongruent, suggesting that at least a subset of cells at this stage is already en route to switch towards the appropriate mesodermal fate.

(E - E') Correlations of the six genes used in the study with (E) *sox2* and (E') *tbxta* were computed and plotted across three stages.

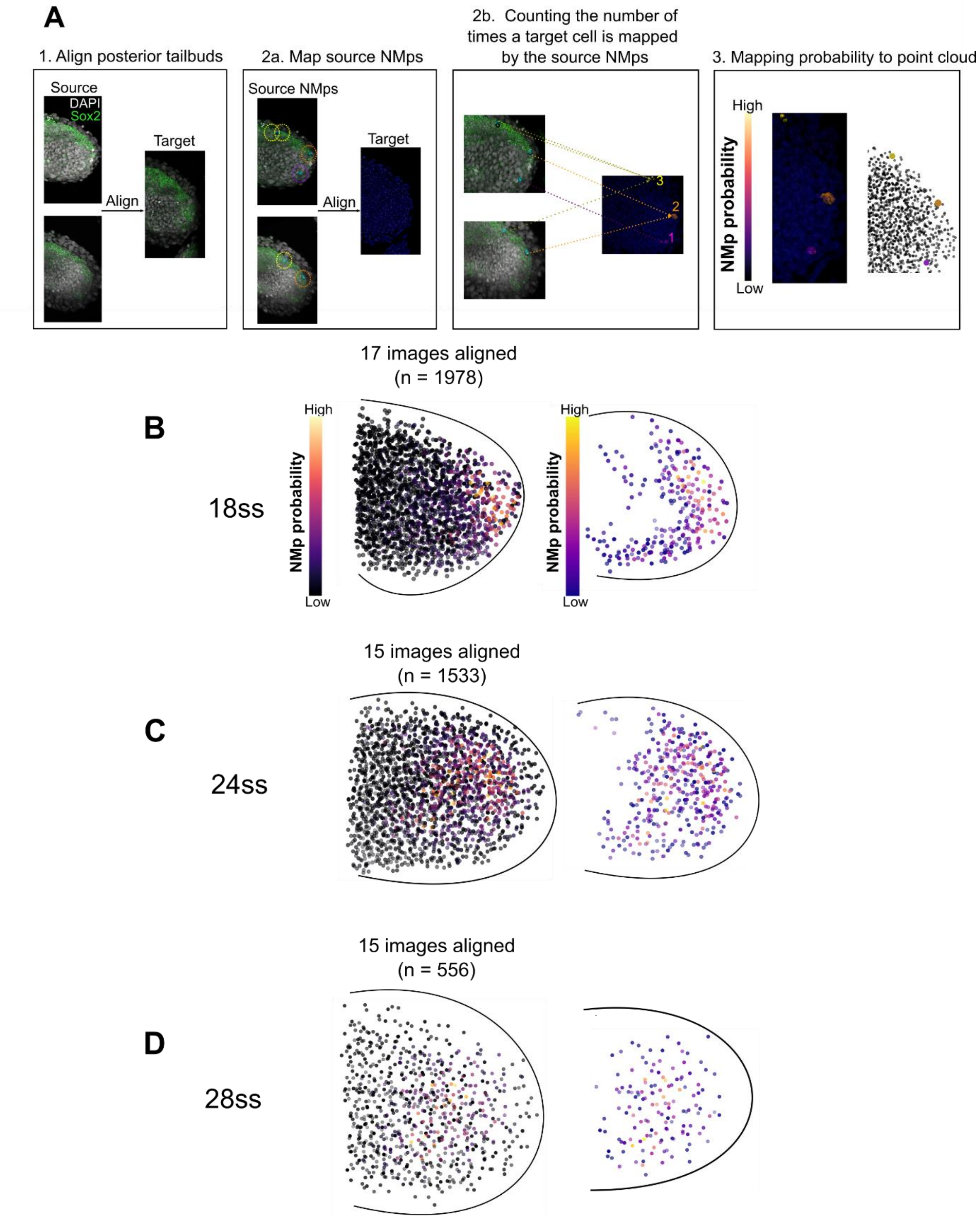


**Figure S14: Construction of an NMp probability map, related to STAR Methods.**

(A) Procedural outline. For each source image that is aligned to the target image, its NMp nuclei has been segmented beforehand in Imaris, and hence we can count the number of times each target cell receives a mapping from a source NMp. Cells with a large count number have a high probability of being an NMP as they are mapped onto very frequently by source NMps. In panels 2b and 3, the yellow surface in the target tailbud has a higher NMp probability than the purple surface as more source NMps map to the yellow surface than the purple surface. Since most target cells are not NMps, the count number is zero for most cells.

(B-D) (Left) NMp probability maps displaying the entire target point cloud. (Right) NMp probability maps displaying only target points with 2 or more mappings from source NMps. The number of tailbud images aligned to the target image, as well as the total number of source NMps used for alignment (n), are displayed on top of each panel.

A

$$\frac{dx}{dt} = (\alpha_1 + \frac{max_1 x^n}{a^n + x^n})(\frac{b^n}{b^n + y^n}) - \lambda_1 x_1$$

$$\frac{dy}{dt} = (\alpha_2 + \frac{max_2 y^n}{c^n + y^n})(\frac{d^n}{d^n + x^n}) - \lambda_2 x_2$$

B

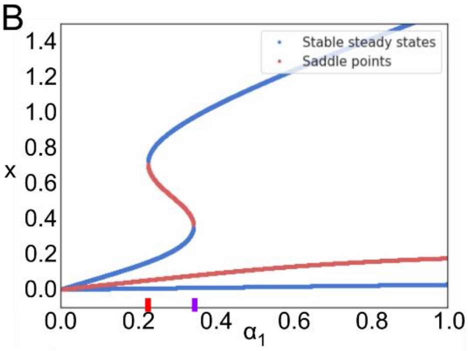


C

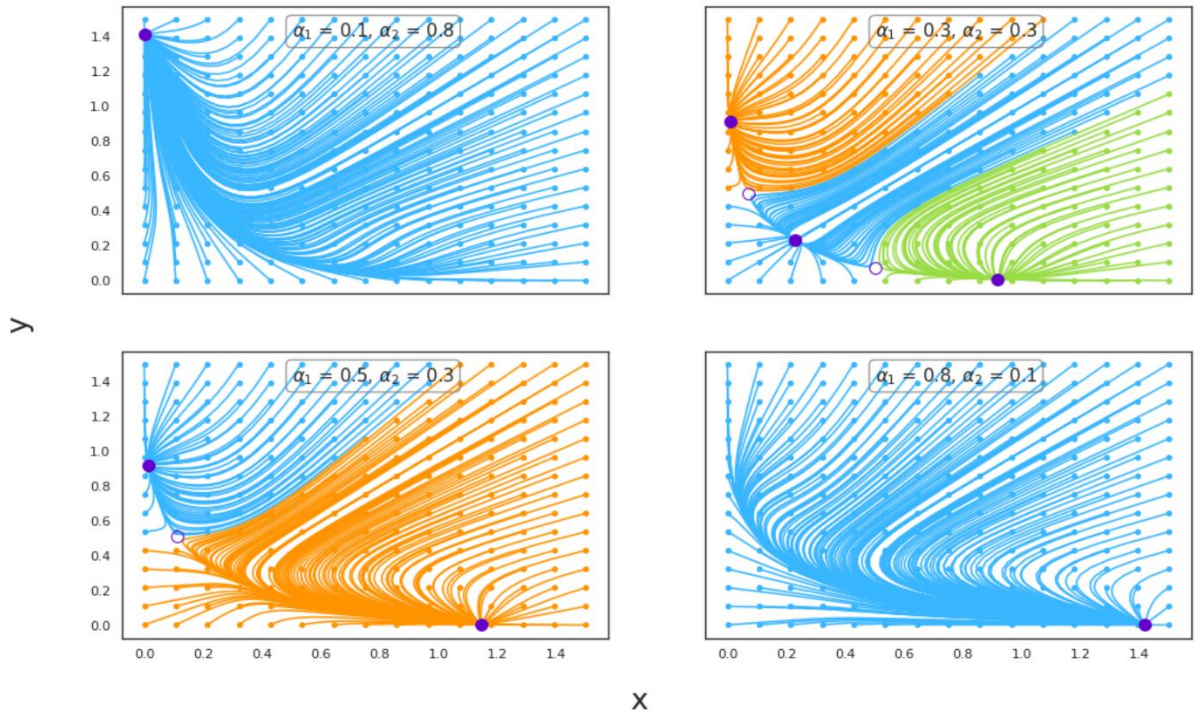


**Figure S15: Deterministic toggle switch, related to STAR Methods.**

(A) The pair of ordinary differential equations (ODEs) used to construct the dynamical model (Equations 3 and 4).

(B) Bifurcation diagram of the toggle switch, showing the solutions of the ODE for gene X as a function of $\alpha_1$, the rate of production of X. The system exhibits bistability (two attractors and one saddle point) at small values of $\alpha_1$. As $\alpha_1$ increases until it reaches a critical threshold (marked as a red line on the x-axis), a sudden qualitative change in the system's behaviour occurs (bifurcation event) - the bifurcation diagram bends over itself, resulting in a total of three attractors and two saddle points. The system remains in this new tristable regime until it reaches another critical threshold (marked as a purple line on the x-axis), where another bifurcation event annihilates the attractor and saddle, returning the system to a bistable regime. Thus, tristability arises when the value of $\alpha_1$ is in a narrow range around $\alpha_2$ ($\alpha_1 \approx \alpha_2$). $\alpha_2 = 0.3$; n = 4; a = c = 0.6; b = d = 0.4; $max_1 = max_2 = 1$; $\lambda_1 = \lambda_2 = 1.25$; Ω = 50.

(C) Phase portraits of the toggle switch at different levels of $\alpha_1$and $\alpha_2$. The stable steady states (attractors) are represented as solid dark blue circles whereas the saddle points are hollow circles. Trajectories are coloured according to the basin of attraction that they reside in. The remaining parameter values are the same as (B).

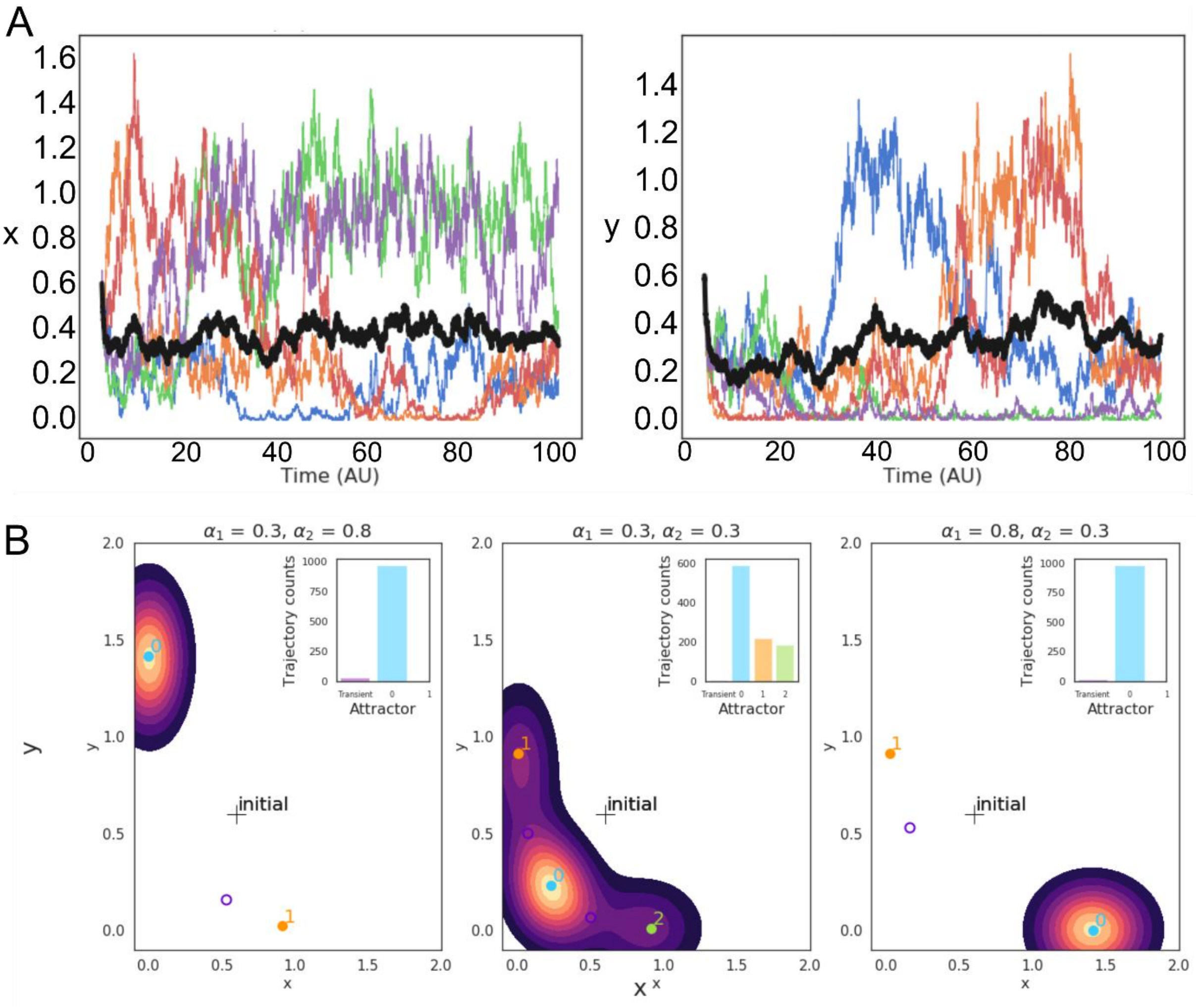


**Figure S16: Stochastic autonomous toggle switch, related to Figure 4.**

(A) Five stochastic trajectories of genes X and Y under the tristable regime ($\alpha_1$= 0.3, $\alpha_2$= 0.3) with identical initial conditions (x = y = 0.6). The mean expression level (bolded black line) is also displayed. The mutual repression between genes X and Y is highlighted in their opposite expression trends - when X is high (low), Y is low (high).

(B) Combined results of 1000 stochastic simulations. The locations of stochastic trajectories at the final 1% time interval of the simulations are shown as density plots imposed onto the state-space. Hotter colours reflect a higher frequency of occurrence. Attractors (filled coloured circles) are labelled numerically. For each state-space plot, the corresponding inset tallies the number of trajectories that have either converged into an attractor or have yet to converge (‘Transient’). In all the simulations, the remaining parameters are the same as Figure S15B.

| Type | Subtype | Gene expression profile consistent with prospective fate | Gene expression profile consistent with Wnt signalling activity |
|---|---|---|---|
| Congruent | | Yes | Yes |
| Incongruent | Compliant | No | Yes |
| | Rebellious | No | No |

**Table S1: Definitions of cell categories, related to Figure 5.**

NMps which possess a gene expression profile that is consistent with its eventual fate and canonical Wnt signalling levels are labelled 'Congruent'. For instance, neural-biased cells located in the neural-fated domain are Congruent cells. Conversely, NMps with a gene expression profile that is inconsistent with their destination fate are labelled 'Incongruent'. These cells can be further subdivided based on their *tcf* (canonical Wnt readout) expression levels. For instance, a mesoderm-biased cell located in the dorsal PW is 'Incongruent'. If its *tcf* expression level is high, the cell would be compliant because high canonical Wnt signalling activity promotes a mesodermal state and this matches its mesodermal-expression profile. Alternatively, it would be 'Rebellious' if its *tcf* level is low as low canonical Wnt activity should promote a neural state and this does not match with its mesodermal expression profile.

| Image Group | Number of images considered for alignment |
|---|---|
| 1) *sox2*, *tbxta*, *tcf* mRNA (STT) | 18ss: 3<br>24ss: 2<br>28ss: 4 |
| 2) *sox2*, *tbxta*, *hes6*, *cdh6* (STHC) | 18ss: 3<br>24ss: 6<br>28ss: 5 |
| 3) *sox2*, *tbxta*, *sp5l*, *cdh6* (STSC) | **18ss: 4**<br>**24ss: 3**<br>28ss: 5 |
| 4) *sox2*, *tbxta*, *tagln3b*, *cdh6* (STTC) | 18ss: 3<br>24ss: 5<br>28ss: 6 |
| 5) *sox2*, *tbxta*, *znf703*, *cdh6* (STZC) | 18ss: 4<br>24ss: 4<br>**28ss: 4** |

**Table S2: Summary of HCR image groups, related to STAR Methods.**

Each image group represents a collection of identical HCR experiments that stain for the genes described. The number of source images considered for the construction of the composite maps at each stage are shown in the right column, although ultimately 1 image from each image group was chosen as the source image to construct the composite map at each stage. Note that 2 target images for the 18ss and 24ss maps (bolded) were chosen from the STSC image group, whilst the target image at 28ss (bolded) was chosen from the STZC image group.

### HCR probe sequences

#### *Sox2* - 16 probe pairs

| B4 split initiators: | Sequence | Spacer |
|---|---|---|
| Initiator_4_odd: | CCTCAACCTACCTCCAAC | AA |
| Initiator_4_even: | TCTCACCATATTCgCTTC | AT |

| Identifier | Sequence |
|---|---|
| 1_B4_*sox2*_odd | cctcaacctacctccaacaaAAGGAAGCGCTCTTAGGTCCTGTTG |
| 2_B4_*sox2*_even | TAGCAGTTGGAGGTAACTTTGCTGGattctcaccatattcgcttc |
| 3_B4_*sox2*_odd | cctcaacctacctccaacaaCGCCGGGGGCTTCAGCTCGGTTTCC |
| 4_B4_*sox2*_even | CCCCGTGCCCCCGGTGTTGGGCTGGattctcaccatattcgcttc |
| 5_B4_*sox2*_odd | cctcaacctacctccaacaaCTGGTTGTTTCCCGAGGAGTTGGTG |
| 6_B4_*sox2*_even | TCTCTTGATGCGGTCCGGGCTGTTTattctcaccatattcgcttc |
| 7_B4_*sox2*_odd | cctcaacctacctccaacaaCGACCACACCATGAAGGCGTTCATG |
| 8_B4_*sox2*_even | CTGTGCCATTTTCCGCCGCTGTCCCattctcaccatattcgcttc |
| 9_B4_*sox2*_odd | cctcaacctacctccaacaaATGAATGGTCGCTTCTCGCTCTCGG |
| 10_B4_*sox2*_even | AGAGCCCGAAGGCGTTTGGCTTCGTattctcaccatattcgcttc |
| 11_B4_*sox2*_odd | cctcaacctacctccaacaaTTGTAATCCGGGTGTTCCTTCATGT |
| 12_B4_*sox2*_even | GTCTTGGTTTTCCTCCGGGGTCTGTattctcaccatattcgcttc |
| 13_B4_*sox2*_odd | cctcaacctacctccaacaaCCGGCCCCTACACCCACGCCGGCGC |
| 14_B4_*sox2*_even | ATCCTCTGGTTGACCCCGGCGCCCAattctcaccatattcgcttc |
| 15_B4_*sox2*_odd | cctcaacctacctccaacaaCAGCCGTTCATATGCGCGTAGCTGT |
| 16_B4_*sox2*_even | TGCATCATGCCGTAGCCTCCGTTGGattctcaccatattcgcttc |
| 17_B4_*sox2*_odd | cctcaacctacctccaacaaGGGTGCTGCGGGTAACCCAGCTGCT |
| 18_B4_*sox2*_even | TGCGCCGTGTTGTGCGCGTTCAAACattctcaccatattcgcttc |
| 19_B4_*sox2*_odd | cctcaacctacctccaacaaATGTCGTAGCGGTGCATGGGCTGCA |
| 20_B4_*sox2*_even | GTCATGGAGTTGTACTGGAGCGCGCattctcaccatattcgcttc |
| 21_B4_*sox2*_odd | cctcaacctacctccaacaaGAGCCGTTCATGTAGGTCTGCGAGT |
| 22_B4_*sox2*_even | TGCGAATAGGACATGCTGTAGGTGGattctcaccatattcgcttc |
| 23_B4_*sox2*_odd | cctcaacctacctccaacaaCCCAGTGTCATTCCCGGCGTGCTTT |
| 24_B4_*sox2*_even | TCGGACTTGACCACCGAGCCCATGGattctcaccatattcgcttc |
| 25_B4_*sox2*_odd | cctcaacctacctccaacaaGTGACCACCGGCGGACTCGAACTAG |
| 26_B4_*sox2*_even | TGTCCGGCCCGGGAATGAGACGACGattctcaccatattcgcttc |
| 27_B4_*sox2*_odd | cctcaacctacctccaacaaATGTCCCGCAGGTCCCCCGTTTGGC |
| 28_B4_*sox2*_even | TCCGCCCCGGGCAGGTACATACTGAattctcaccatattcgcttc |
| 29_B4_*sox2*_odd | cctcaacctacctccaacaaCTGCTCTGAGCGCTCTGGTCCTGCA |
| 30_B4_*sox2*_even | CTCTGGTAATGTTGGGACATGTGCAattctcaccatattcgcttc |
| 31_B4_*sox2*_odd | cctcaacctacctccaacaaTTAATCGTCGTACCGGGCACAGGTG |
| 32_B4_*sox2*_even | TACATATGCGATAAGGGAATCGTGCattctcaccatattcgcttc |

#### *Tbxta* - 26 probe pairs

| **B1 split initiators:** | **Sequence** | **Spacer** |
|---|---|---|
| Initiator_1_odd: | gAggAgggCAgCAAACgg | AA |
| Initiator_1_even: | gAAgAgTCTTCCTTTACg | TA |

| **Identifier** | **Sequence** |
|---|---|
| 1_B1_*tbxta*_odd | gaggagggcagcaaacggaaACTGTTGCTTTGACAGCGGGAATTC |
| 2_B1_*tbxta*_even | CGACGATCCTACTAAATCCCGTTGGtagaagagtcttcctttacg |
| 3_B1_*tbxta*_odd | gaggagggcagcaaacggaaGGACTTGAGGCAGACATATTTCCGA |
| 4_B1_*tbxta*_even | CTAAGGAGATGATCCAGGCGCTGGTtagaagagtcttcctttacg |
| 5_B1_*tbxta*_odd | gaggagggcagcaaacggaaCCCTTCTGAAATTCGCTCTCCACGG |
| 6_B1_*tbxta*_even | CGCTCGGACGCGTCCCCTTTCTCGCtagaagagtcttcctttacg |
| 7_B1_*tbxta*_odd | gaggagggcagcaaacggaaCTCCGCGTCTTCAAGCGAAAGTTTA |
| 8_B1_*tbxta*_even | TTGGTGAGCTCTTTAAATTTGGTCCtagaagagtcttcctttacg |
| 9_B1_*tbxta*_odd | gaggagggcagcaaacggaaCACGGGAAACATTCGTCTCCCAGTC |
| 10_B1_*tbxta*_even | GTCGAGACCGGTGACACTGGCTCTGtagaagagtcttcctttacg |
| 11_B1_*tbxta*_odd | gaggagggcagcaaacggaaCAGCAGGACCGAGTACATTGCATTA |
| 12_B1_*tbxta*_even | CCGATTATTATCGGCCGCCACAAAAtagaagagtcttcctttacg |
| 13_B1_*tbxta*_odd | gaggagggcagcaaacggaaGGGCACCCATTCACCGTTCACGTAT |
| 14_B1_*tbxta*_even | CGGGCTTTGGGGTTCGGGTTTCCCAtagaagagtcttcctttacg |
| 15_B1_*tbxta*_odd | gaggagggcagcaaacggaaTGAGTCCGGGTGGATGTAGACGCAG |
| 16_B1_*tbxta*_even | TTTCATCCAGTGCGCGCCGAAGTTGtagaagagtcttcctttacg |
| 17_B1_*tbxta*_odd | gaggagggcagcaaacggaaTTTGACTTTGCTGAAAGATACGGGT |
| 18_B1_*tbxta*_even | TAACATAATCTGTCCTCCTCCGTTGtagaagagtcttcctttacg |
| 19_B1_*tbxta*_odd | gaggagggcagcaaacggaaGACTTTCACGATGTGTATCCTGGGT |
| 20_B1_*tbxta*_even | ACTGCTGATCATTTTCTGAATCCCAtagaagagtcttcctttacg |
| 21_B1_*tbxta*_odd | gaggagggcagcaaacggaaGTGACTGCAATAAACTGTGTCTCAG |
| 22_B1_*tbxta*_even | GTGTTTGATTTTCAGAGCGGTAATCtagaagagtcttcctttacg |
| 23_B1_*tbxta*_odd | gaggagggcagcaaacggaaACTTCTCTCTTTGGCATCGAGGAAA |
| 24_B1_*tbxta*_even | GCTGTGGTCTGGGACTTCCTTGTGGtagaagagtcttcctttacg |
| 25_B1_*tbxta*_odd | gaggagggcagcaaacggaaTGAATATCCAGATTGCTGGTTGTCA |
| 26_B1_*tbxta*_even | ACTGGGCAGGAACCAGCCACCGAGTtagaagagtcttcctttacg |
| 27_B1_*tbxta*_odd | gaggagggcagcaaacggaaGCTGCTGCTGGGGCCCATGGGGCCG |
| 28_B1_*tbxta*_even | AACAGGGGCCCCATTGAACTGAGGAtagaagagtcttcctttacg |
| 29_B1_*tbxta*_odd | gaggagggcagcaaacggaaTCTCTCACAGTACGAACCCGAGGAG |
| 30_B1_*tbxta*_even | AGCTCTGTGGTTCCTCAAGCTGGAGtagaagagtcttcctttacg |
| 31_B1_*tbxta*_odd | gaggagggcagcaaacggaaGTGGGAGTAATGGCTGGGATATGGA |
| 32_B1_*tbxta*_even | CATGTAGTTATTGGTGGTAGTGCTGtagaagagtcttcctttacg |
| 33_B1_*tbxta*_odd | gaggagggcagcaaacggaaGAGACGCAAGACTTCCGGAAGAGTT |
| 34_B1_*tbxta*_even | GGATCTGCAGGGCTGACCAGCTGTCtagaagagtcttcctttacg |
| 35_B1_*tbxta*_odd | gaggagggcagcaaacggaaCCAGGGTTCCCATCCCGCTGGAGTT |
| 36_B1_*tbxta*_even | TGTTGGAGGTAGTGTTTGTGGTGTGtagaagagtcttcctttacg |
| 37_B1_*tbxta*_odd | gaggagggcagcaaacggaaCTGACCACAGACTTGGGTACTGACT |
| 38_B1_*tbxta*_even | CTGATGGGGTGAGAGTCGTCCCTGCtagaagagtcttcctttacg |
| 39_B1_*tbxta*_odd | gaggagggcagcaaacggaaCACCTGTAATGGAGCCCGATGCTGA |
| 40_B1_*tbxta*_even | AACCGCGTAGGAACTGAGATGTCAGtagaagagtcttcctttacg |
| 41_B1_*tbxta*_odd | gaggagggcagcaaacggaaAGGTCAGACCCGAGTAGGACATCGA |
| 42_B1_*tbxta*_even | AGGAGGGAGAGGACACAGGCAGCGAtagaagagtcttcctttacg |
| 43_B1_*tbxta*_odd | gaggagggcagcaaacggaaCCTCGCTTAGGCCTGGATCGTACAT |
| 44_B1_*tbxta*_even | TCTCGAACTGGGCATCTCCAACGCCtagaagagtcttcctttacg |
| 45_B1_*tbxta*_odd | gaggagggcagcaaacggaaATGATGCTGTGAGCCGGGCGATGGA |
| 46_B1_*tbxta*_even | CTCAGTAGCTCTGAGCCACAGGCGCtagaagagtcttcctttacg |
| 47_B1_*tbxta*_odd | gaggagggcagcaaacggaaAGCATCAGTCCTTAAATGTGAAGCG |
| 48_B1_*tbxta*_even | CTGAAGCCAAGATCAAGTCCATAACtagaagagtcttcctttacg |
| 49_B1_*tbxta*_odd | gaggagggcagcaaacggaaCAACCCGTTTTCTGATTGTCAAATC |

| | |
|---|---|
| 50_B1_*tbxta*_even | CACCCTCAAGTTGTTCCAAACTTTAtagaagagtcttcctttacg |
| 51_B1_*tbxta*_odd | gaggagggcagcaaacggaaTCTTCTGTGATACAATGAAACCGGA |
| 52_B1_*tbxta*_even | CAGCAAAGTCTGTCTTTCTCTCGTTtagaagagtcttcctttacg |
